# Epidermal loss of PRMT5 leads to the emergence of an atypical basal keratinocyte-like cell population and defective skin stratification

**DOI:** 10.1101/2024.11.08.620904

**Authors:** Nicole Recka, Andrean Simons, Robert A. Cornell, Eric Van Otterloo

## Abstract

During skin development, ectoderm-derived cells undergo precisely coordinated proliferation, differentiation, and adhesion to yield stratified epidermis. Disruptions in these processes can result in congenital anomalies including ectodermal dysplasia and harlequin ichthyosis. Protein Arginine Methyl Transferase 5 (PRMT5)—an enzyme responsible for methylating arginine residues in histones and other proteins—maintains progenitor status in germ and limb bud cells. Similarly, *in vitro* evidence suggests that PRMT5 prevents differentiation of basal keratinocytes, leading us to hypothesize that PRMT5 preserves the stem-cell phenotype of keratinocytes *in vivo*. To test this possibility, we generated conditional knockout (cKO) mice lacking *Prmt5* in early ectoderm (E7.5), impacting the entire epidermis. *Prmt5* cKOs exhibited gross skin defects, compromised skin barrier function, and reduced postnatal viability. Histological analyses revealed significant defects in epidermal stratification, without alterations in apoptosis or proliferation. Single-cell RNA and ATAC-seq analysis identified an atypical population of basal keratinocyte-like cells in *Prmt5* cKOs, that exhibited a senescence-like program, characterized by increased *Cdkn1a* (p21), elevated senescence-associated secretory phenotype (SASP) molecules (*Igfbp2*), and decreased developmental transcription factor (*Trp63*) expression. Our findings suggest that PRMT5 prevents basal keratinocyte senescence by repressing *Cdkn1a*, shedding light on the epigenetic regulation of basal keratinocyte maintenance and senescence in congenital skin disorders.

## INTRODUCTION

Skin is a rapidly regenerating organ that serves as the body’s primary defense against the range of external threats it faces daily. Well-functioning skin relies on the proper development of the epidermis to create a water-resistant barrier, crucial for regulating an organism’s internal body temperature, preventing dehydration, and protecting against invading pathogens. Deviations from proper epidermal development and homeostasis can result in various skin conditions, ranging from mild cases of psoriasis and eczema to severe congenital anomalies such as harlequin ichthyosis and epidermolysis bullosa (Coulombe et al., 1991, Dale et al., 1990, Sharma, 2001, Weinstein and Frost, 1968). To improve clinicians’ capacity for genetic diagnosis and targeted treatments of skin-related pathologies, a deeper comprehension of the molecular drivers underlying normal skin development is essential.

The epidermis originates from a single layer of surface ectoderm, driven towards epidermal commitment by a sequence of BMP and Notch signaling events (Wilson et al., 2001). Within the newly formed basal layer, cells possess the capacity for division either in a plane parallel to the underlying dermis to populate and perpetuate the basal layer, or in a plane perpendicular to the dermis, leading to stratification and formation of the spinous layer (Lavker and Sun, 1983). Spinous cells undergo further stratification to create the granular layer whose cells divide a final time to create the outermost cornified layer (Steven et al., 1990). As cells ascend through these layers, they progressively withdraw from the normal cell cycle, culminating in a densely packed cornified husk of denuclearized cells that establishes a water-resistant barrier (Sun and Green, 1976). Cells in each layer of the epidermis express a unique combination of transcription factors, morphogens, structural keratins, and other functional proteins to create a distinct molecular environment. While there has been success in identifying and modulating essential genes within this genetic network (e.g., *Tp63*, *Irf6*, *Notch*) (Fuchs, 2007), the regulation of skin development by enzymes that remodel chromatin remains to be fully elucidated.

Chromatin is modified by post-translational modifications of histones, including acetylation, methylation, and ubiquitination (Bannister and Kouzarides, 2011). Among such enzymes participating in these modifications are the protein arginine methyltransferases (PRMTs) which catalyze mono and di-methylation of arginine residues in histones and other proteins (Stopa et al., 2015). One family member, PRMT5, mediates symmetric di-methylation of arginines in histones (Pal et al., 2004), transcription factors (Hu et al., 2015), RNA splicing enzymes (Gonsalvez et al., 2007), and other proteins (Hou et al., 2008). PRMT5 also appears to maintain a progenitor, stem-cell fate in germ cells and cells of the developing limb, as conditional deletion in premeiotic male germ cells or limb bud mesenchyme leads to a reproductive defect and syndactyly, respectively (Norrie et al., 2016, Wang et al., 2015). Previous studies have indicated that PRMT5 similarly inhibits *in vitro* keratinocyte differentiation (Kanade and Eckert, 2012), however this function has not been confirmed *in vivo*. In this study, we investigated the necessity of PRMT5 during *in vivo* skin development using an epidermal conditional (cKO) mouse model.

Our findings revealed that PRMT5 loss in the epidermis leads to severe defects in stratification, improper barrier formation, and perinatal lethality. Single-cell RNA and ATAC sequencing (scRNA/ATAC-seq) of skin from *Prmt5* cKO mice revealed an abnormal population of keratinocytes characterized by a senescence-like program. These cells exhibited marked upregulation of the senescence driver *Cdkn1a* (*p21*), overexpression of the senescence-associated secretory phenotype (SASP) molecule *Igfbp2*, and decreased expression of *Trp63*, a key transcription factor in skin development. These findings highlight the critical role of PRMT5 in proper epidermal development and may provide insight into the role of methyltransferases in skin-related pathologies.

## RESULTS

### PRMT5 is expressed throughout epidermal development and can be conditionally deleted with the ectodermal-CRE, *Crect*

To complement an earlier analysis of PRMT5 expression during mouse development (Tee et al., 2010), we performed immunofluorescence using a PRMT5-specific antibody at key stages of epidermal development (De Falco et al., 2014), focusing on facial epidermis. At the onset of epidermal specification (embryonic day 13.5, or E13.5), anti-PRMT5 immunoreactivity was detected throughout the largely single-layered epidermis and the loosely packed dermis (Fig. 1A). At initial stages of epidermal stratification (E15.5), anti-PRMT5 immunoreactivity remained widespread in the dermis and both basal and suprabasal layers of the epidermis (Fig. 1B). However, anti-PRMT5 localization varied between epidermal and dermal layers, with cytoplasmic enrichment noted in the epidermis and nuclear enrichment in the dermis (Fig. 1B). At later stages of epidermal stratification and maturation (E17.5), anti-PRMT5 immunoreactivity persisted throughout the epidermis and dermis, with immunoreactivity enriched in the basal layer relative to more superficial layers of epidermis (Fig. 1C). Thus, we have confirmed that PRMT5 is expressed at key stages of epidermal development.

**Figure 1:**
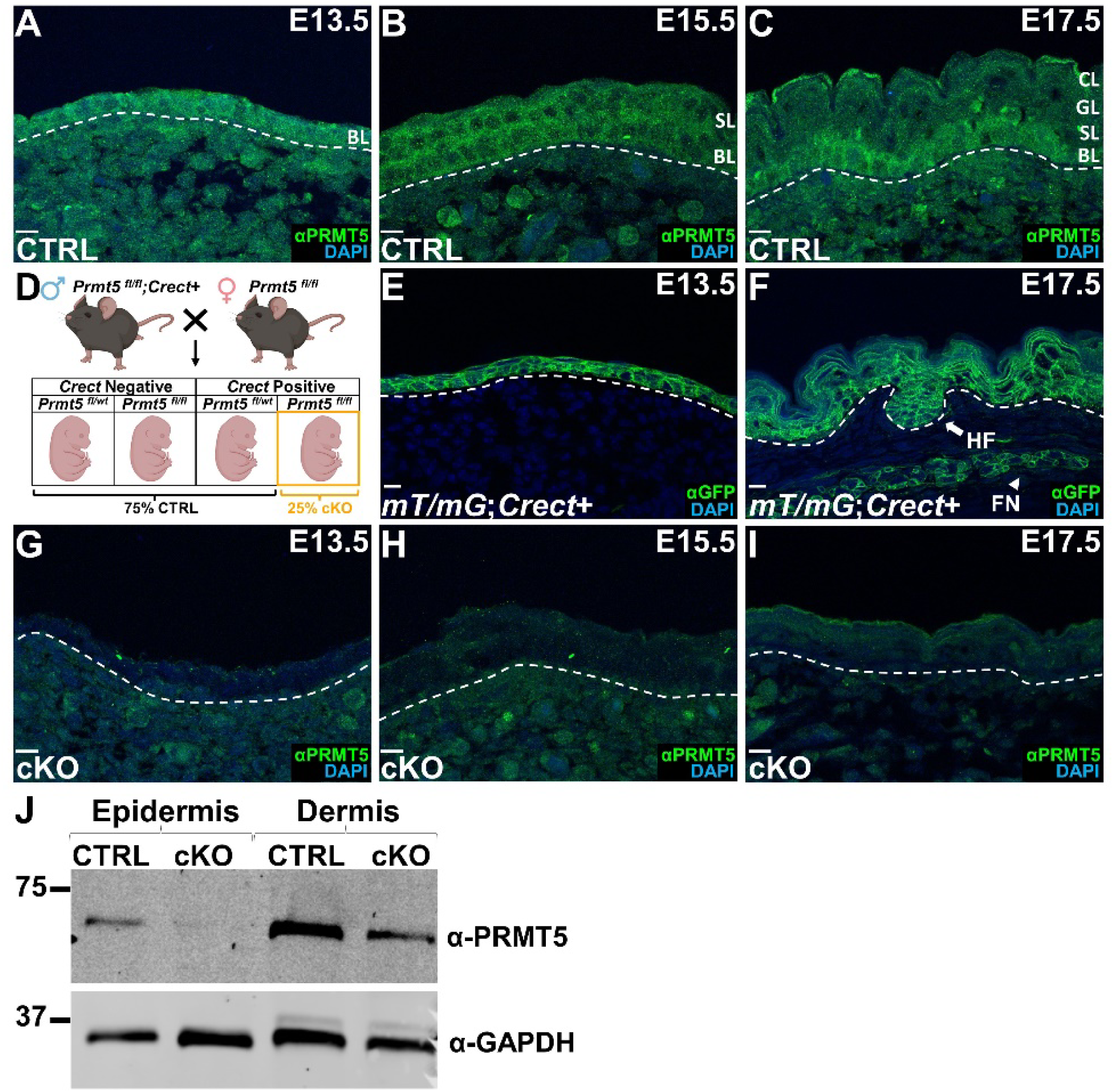
PRMT5 is expressed throughout epidermal development and can be conditionally deleted using *Crect*. (**A-C**) α-PRMT5 sectional immunofluorescence in a control (i.e., Cre negative, CTRL) mouse embryo at key timepoints of epidermal development, including E13.5 (A), E15.5 (B), and E17.5 (C). (**D**) Schematic depicting the breeding scheme used in this study. Briefly, a *Prmt5 fl/wt;Crect+* male is bred with a *Prmt5 fl/fl* female, producing ∼25% *Prmt5 fl/fl;Crect+* (referred to as ‘cKO’) embryos along with additional genotypes (collectively referred to as ‘CTRL’ embryos). (**E-F**) α-eGFP sectional immunofluorescence in an E13.5 (E) or E17.5 (F) embryo containing both *Crect* and the *mT/mG* reporter allele, allowing detection of CRE-mediated recombination. (**G-I**) α-PRMT5 sectional immunofluorescence in a cKO mouse embryo at timepoints matched to those of their respective sibling CTRL (A-C)—E13.5 (G), E15.5 (H), and E17.5 (I). (**J**) Western blot analysis with an α-PRMT5 antibody detecting PRMT5 expression in protein extracts isolated from E18.5 epidermal or dermal tissue in a CTRL or cKO embryo. GAPDH serves as a loading control. Note, sections shown are the craniofacial epidermis, lateral to the oral cavity, in a coronal plane. White dashed lines demarcate the boundary of the dermis and epidermis. Acronyms include basal layer (BL), spinous layer (SL), granular layer (GL), cornified layer (CL), hair follicle (HF), and facial nerve (FN). Nuclei are counterstained with DAPI. All sectional immunofluorescence images are five Z-stacks compiled with Max Intensity in FIJI. Scale bar = 10µm.

As *Prmt5*-null embryos are early embryonic lethal (∼E6.5) (Tee et al., 2010), precluding assessment of PRMT5 function in the epidermis, we coupled a *Prmt5* floxed allele (Norrie et al., 2016) with an ectodermally-expressed CRE, *Crect* (Reid et al., 2011), to conditionally delete *Prmt5* from the ectoderm (*Prmt5* cKO) and its derivatives (Fig. 1D). Crossing mice with *Crect*-driven CRE to those harboring recombination reporter alleles (mT/mG, (Muzumdar et al., 2007) or r26r, (Soriano, 1999)) yielded embryos with GFP expression or ß-galactosidase staining, respectively, within ectoderm-derived tissues (Fig. 1E, F, Supp. Fig. 1), confirming efficacy and specificity of *Crect*-driven CRE. Relevant to this study, *Crect* is expressed prior to the onset of skin development in the head and trunk, resulting in reporter expression throughout all early (i.e., E13.5, Fig. 1E) and late (i.e., E17.5, Fig. 1F) stage layers of the epidermis and oral epithelium (Supp. Fig. 1). Note, given the rare occurrence of ectopic CRE expression with the *Crect* allele (Schock et al., 2017), each breeder male’s *Crect* expression pattern was verified using the mTmG reporter, with those exhibiting ectopic expression excluded (Supp. Fig. 1). In the head, we also noted recombination in the presumed ectoderm-derived facial nerves at E17.5, as identified by localization of α-GFP and α-neurofilament immunostaining (Fig. 1F, Supp. Fig. 1).

Consistent with the reporter alleles, α-PRMT5 antibody immunofluorescence on tissue sections of control (CTRL) embryos (Fig. 1A-C) and *Prmt5* cKOs (Fig. 1G-I) revealed epidermal-specific loss of PRMT5 in the latter. While expression in the dermis was largely unaffected at earlier stages (i.e., E13.5 and E15.5, Fig. 1G, H) in *Prmt5* cKOs, we noted an apparent reduction at later stages (i.e., E17.5, Fig. 1I). Consistent with immunofluorescence, Western blot analysis of protein isolated from back skin of E18.5 CTRL or *Prmt5* cKO embryos revealed a near complete loss of PRMT5 protein in the epidermis of *Prmt5* cKOs (Fig. 1J). In contrast, PRMT5 expression persisted in the dermis of both CTRL and *Prmt5* cKO embryos, although we again noted a mild reduction in the dermis of *Prmt5* cKOs, relative to CTRL at this late stage (Fig. 1J). Given the lack of detected recombination in the dermis, this late-stage reduction in the dermis may result indirectly from defects in the epidermis. In sum, we found that PRMT5 is expressed throughout embryonic epidermal development in controls and this expression is absent in the epidermis of *Prmt5* cKOs.

### Epithelial-specific loss of PRMT5 results in perinatal lethality associated with gross epidermal defects

Having characterized PRMT5 expression in the skin and confirmed its targeted deletion in the epidermis, we next assessed how loss of PRMT5 impacts embryonic and postnatal development. Genotyping of embryos at E15.5 (n=185) and E18.5 (n=128) and of mice at weaning stage (i.e., postnatal day (P) 21 or P21) revealed the correct number (i.e., expected by Mendelian frequencies) of *Prmt5* cKO embryos at E15.5 and E18.5, but significantly fewer than expected at P21 (n=112, p < 0.0001) (Fig. 2A), suggesting perinatal lethality of *Prmt5* cKO animals.

**Figure 2:**
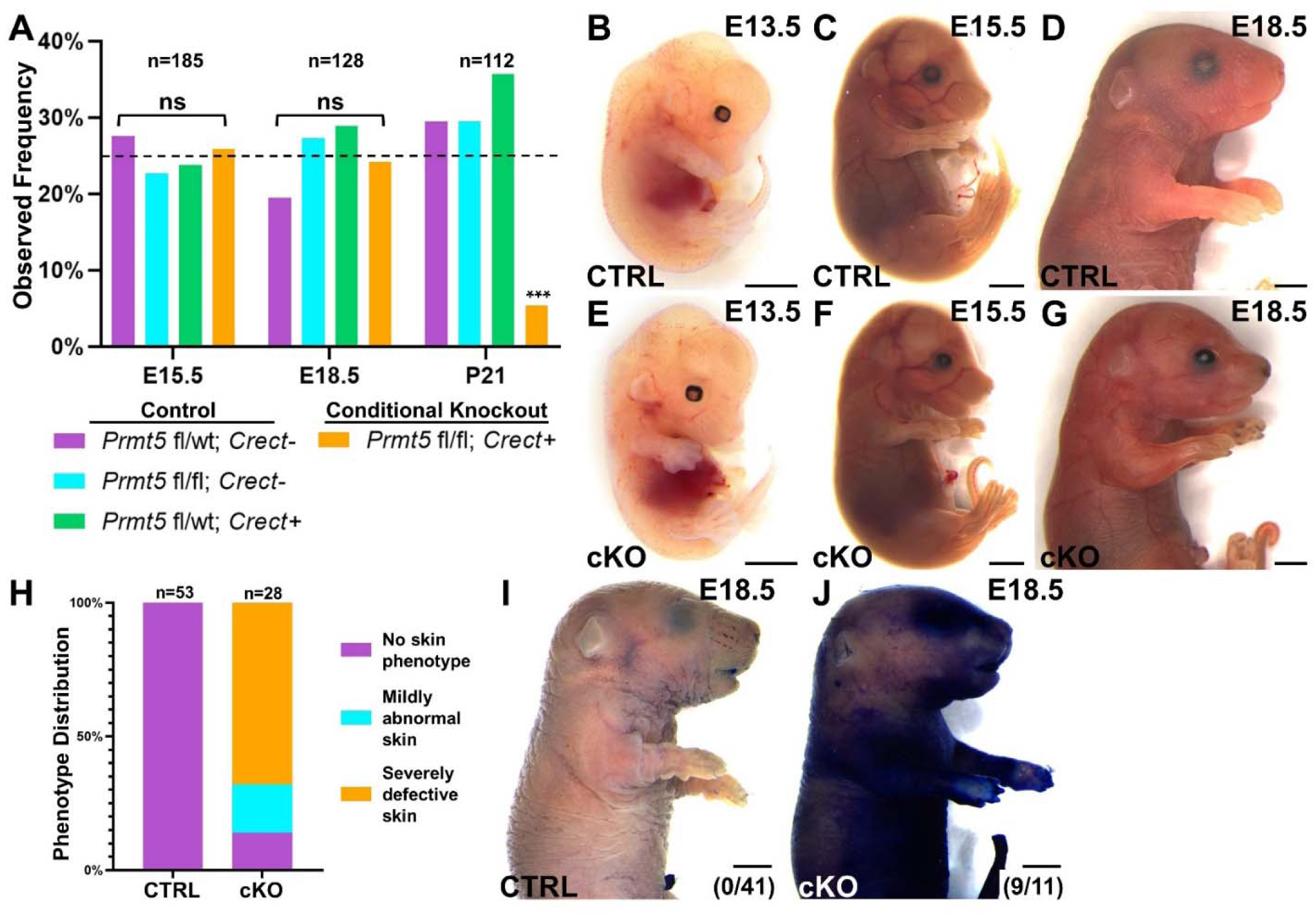
Epithelial-specific loss of PRMT5 results in lethality, gross skin defects, and a compromised epidermal barrier. (**A**) A histogram summarizing the observed frequency (Y-axis) of the possible genotypes, generated from the experimental cross, at different embryonic (e.g., E15.5, E18.5) and postnatal (e.g., P21) stages (X-axis). The expected frequency for each genotype (i.e., 25%) is indicated by the dashed line. Note, ‘n =’ refers to the total number of embryos/pups scored at the indicated stage, ns refers to ‘not significant’, and *** = p < 0.0001, Likelihood Ratio (RStudio package XNomial). (**B-I**) Brightfield lateral view images of a CTRL (B-D, I) or cKO (E-G, J) embryo at E13.5 (B, E), E15.5 (C, F), and E18.5 (D, I-J) stages. Ventral is to the right. Embryos in I and J have also been stained with toluidine blue as an assessment of skin permeability, with the number of embryos displaying permeable skin quantified in the bottom right corner (I-J). (**H**) A bar graph qualitatively summarizing no (purple), mild (cyan), or severe (orange) skin defects in CTRL and cKO samples. Note, gross anatomy images underwent post-acquisition white balancing using GIMP. Scale bar = 2mm.

To assess embryonic phenotypes preceding perinatal lethality, we examined CTRL and *Prmt5* cKO embryos at various developmental timepoints (Fig. 2B-G). At E13.5, associated with the onset of epidermal specification, *Prmt5* cKO mice looked indistinguishable from their CTRL littermates (Fig. 2B, E). However, at E15.5, associated with early stages of epidermal stratification, subtle alterations in the gross appearance of the skin and a distinctive kinked tail were apparent in *Prmt5* cKOs, relative to CTRLs (Fig. 2C, F). By E18.5, associated with the completion of skin development, these alterations became readily apparent with over 85% of *Prmt5* cKO exhibiting obvious skin defects (Fig. 2D, G). Of these, the majority (∼68%) were classified as ‘severe’ and displayed taut and shiny skin, lacking the normal topology of CTRL skin (Fig. 2D, G, H). While a subset (∼18%) showed similar, but less severe features (Fig. 2H, Supp. Fig. 2). Additional epidermal-associated defects were noted in *Prmt5* cKOs, relative to CTRLs, although they displayed variable penetrance and expressivity. Defects included fused digits, open eyelids, missing whiskers, a kinked tail, and oral blistering (Supp. Fig. 2).

Finally, given the observable skin defects and perinatal lethality—often associated with a defective skin barrier—skin permeability was assessed using a toluidine blue assay (Hardman et al., 1998) in E18.5 CTRL and *Prmt5* cKO embryos. While all CTRL embryos (n = 41/41) resisted staining by toluidine blue (Fig. 2I), the majority of *Prmt5* cKO embryos (n = 9/11) readily retained toluidine blue (Fig. 2J), indicative of a functional defect in skin maturation in *Prmt5* cKOs. In sum, phenotypic analyses indicated that PRMT5 function in the epidermis is required for proper skin development, formation of a permeability barrier, and perinatal survival.

### PRMT5-deficient epidermis fails to undergo proper stratification but retains normal proliferation and survival

Having established that loss of *Prmt5* expression results in a severe skin defect at a gross level, we next used histology and immunofluorescence (IF) to compare CTRL and *Prmt5* cKO epidermis at a cellular level. Consistent with the mild defects observed at early stages of skin development, Hematoxylin and Eosin (H&E) staining of E15.5 CTRL and *Prmt5* cKO sections revealed little difference in epidermal organization (Fig. 3A, B). In contrast, H&E staining of E18.5 CTRL sections revealed appropriate stratification (i.e., basal, spinous, granular, and cornified layers all evident) but severely hypoplastic epidermis in *Prmt5* cKO sections, characterized by an apparent absence of spinous or granular layers, a halo of sparse cornified layer, and a reduced basal layer (Fig. 3C-D).

**Figure 3:**
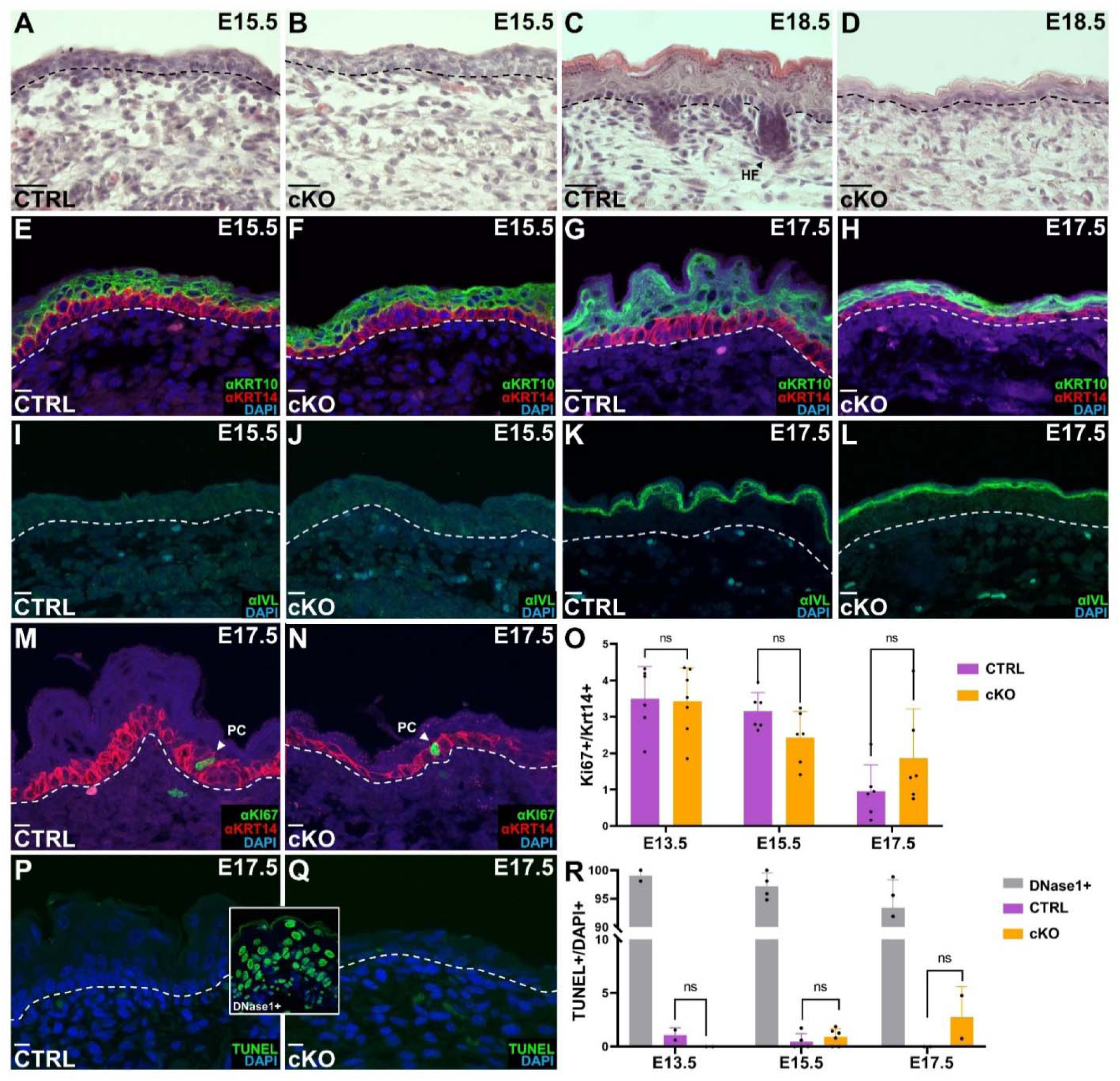
PRMT5-deficient skin fails to undergo proper stratification but retains normal proliferation and death. (**A-H**) Coronal sections from early (E15.5) (A-B, E-F, I-J) or late (E17.5, E18.5) (G-H, C-D, K-L) stage CTRL (A, C, E, G, I, K) or cKO (B, D, F, H, J, L) embryonic heads, processed by Hematoxylin & Eosin (H&E) staining (A-D), α-KRT14 and α-KRT10 immunofluorescence (E-H), or α-IVL immunofluorescence (I-L). (**M, N**) An E17.5 CTRL (M) or cKO (N) section stained by immunofluorescence for α-KRT14 and α-Ki67 to monitor basal cell proliferation. (**O**) Histograms summarizing Ki67+ cells within the epidermis of CTRL and cKO embryos at three different embryonic stages. test. Note, each dot signifies one embryo per genotype and ns refers to ‘not significant’ as determined by a student t-test. (**P, Q**) An E17.5 CTRL (P) or cKO (Q) section processed for TUNEL staining to monitor cell death. Sections were also pre-treated with DNase1 to generate TUNEL+ cells as a positive control. (**R**) Histograms summarizing TUNEL+ cells within the epidermis of DNase1 treated, CTRL, and cKO embryos at three different embryonic stages. Note, sections shown are the craniofacial epidermis, lateral to the oral cavity, in a coronal plane. White dashed lines demarcate the boundary of the dermis and epidermis. Acronyms include hair follicle (HF) and proliferating cell (PC). Nuclei are counterstained with DAPI. All sectional immunofluorescence images are five Z-stacks compiled with ‘Max Intensity’ in FIJI. Scale bar in A-D = 25µm. Scale bar in E-N, P-Q = 10µm.

Given evidence from *in vitro* studies that PRMT5 prevents differentiation of keratinocytes (Kanade and Eckert, 2012), we conducted IF with antibodies recognizing distinct layers of the stratified epithelium. We first used α-KRT14, α-KRT10, α-IVL, and α-KRT6 antibodies to recognize the basal, suprabasal, differentiated, and periderm layers, respectively. At E15.5, *Prmt5* cKOs displayed qualitatively normal KRT14, KRT10, IVL, and KRT6 expression and epidermal morphology, supporting normal differentiation until this timepoint (Fig. 3E, F, I, J, Supp. Fig. 3). By contrast, at E17.5, *Prmt5* cKOs had a hypoplastic epidermis—like that observed in E18.5 H&E sections (Fig. 3C, D)—and reduced albeit present, domains of KRT14, KRT10, and IVL expression relative to in CTRLs (Fig. 3G, H, K, L). In addition, KRT14-expressing basal cells appeared smaller and flatter in *Prmt5* cKOs relative to in CTRLs (Fig. 3G, H). Note, as the periderm is largely absent by E16.5 (Richardson et al., 2014), we did not assess KRT6 expression at this stage. Second, we used α-COL4, α-VCL, and α-CLDN1 antibodies to assess structural proteins present in the basement membrane, adherens junctions, and tight junctions, respectively (Supp. Fig. 3). At E15.5, both *Prmt5* cKOs and CTRLs displayed similar COL4 expression within the basement membrane of basal keratinocytes, VCL expression at cell-cell junctions throughout basal and suprabasal layers, and CLDN1 expression at tight junctions throughout the suprabasal layers (Supp. Fig. 3).

Finally, given the reduced epidermal thickness in *Prmt5* cKOs relative to in CTRLs, we quantified cell proliferation and death throughout skin development (i.e., at E13.5, E15.5, E17.5). α-Ki67 IF revealed comparable ratios of proliferating basal keratinocytes (i.e., KI67+/KRT14+ double positive/ total KRT14+) in *Prmt5* cKOs and in CTRLs across timepoints, although at E17.5 the ratio trended upwards in *Prmt5* cKOs compared to CTRLs (Fig. 3M-O). Similarly, TUNEL staining revealed similar rates of apoptosis between genotypes, across timepoints (Fig. 3P-R). In sum, histology and IF identified that loss of PRMT5 from the surface epithelium results in severe skin hypoplasia and stratification defects, associated with abnormal keratinocyte morphology at late gestational stages. However, these defects do not appear to be correlated with dramatic changes in keratinocyte differentiation, proliferation, survival, or cell-cell/cell-matrix contacts.

### Single cell multiome sequencing identifies reduced epidermal populations, concomitant with an expanded novel keratinocyte-like cell type, in *Prmt5* cKO skin

To determine the impact of *Prmt5* epidermal deletion on the cellular, transcriptomic and chromatin-accessibility landscapes of individual cells within the developing skin, we isolated full-thickness dorsal trunk skin from E15.5 CTRL and *Prmt5* cKO embryos, dissociated the skin into a single cell suspension, isolated nuclei, and generated paired single-cell accessibility (scATAC-seq) and expression (scRNA-seq) libraries using the 10X Genomics Chromium system (Fig. 4A). As E15.5 is associated with only mild phenotypic changes, we predicted that this timepoint would reveal early cellular and molecular changes caused by *Prmt5* deletion with minimal secondary effects. To assure biological relevance, for each sample (i.e., CTRL and *Prmt5* cKO) we pooled three embryos from matched litters. After quality control, our dataset contained 17,696 and 14,319 cells for CTRL and *Prmt5* cKO samples, respectively (Fig. 4B, Supp. Fig. 4).

**Figure 4:**
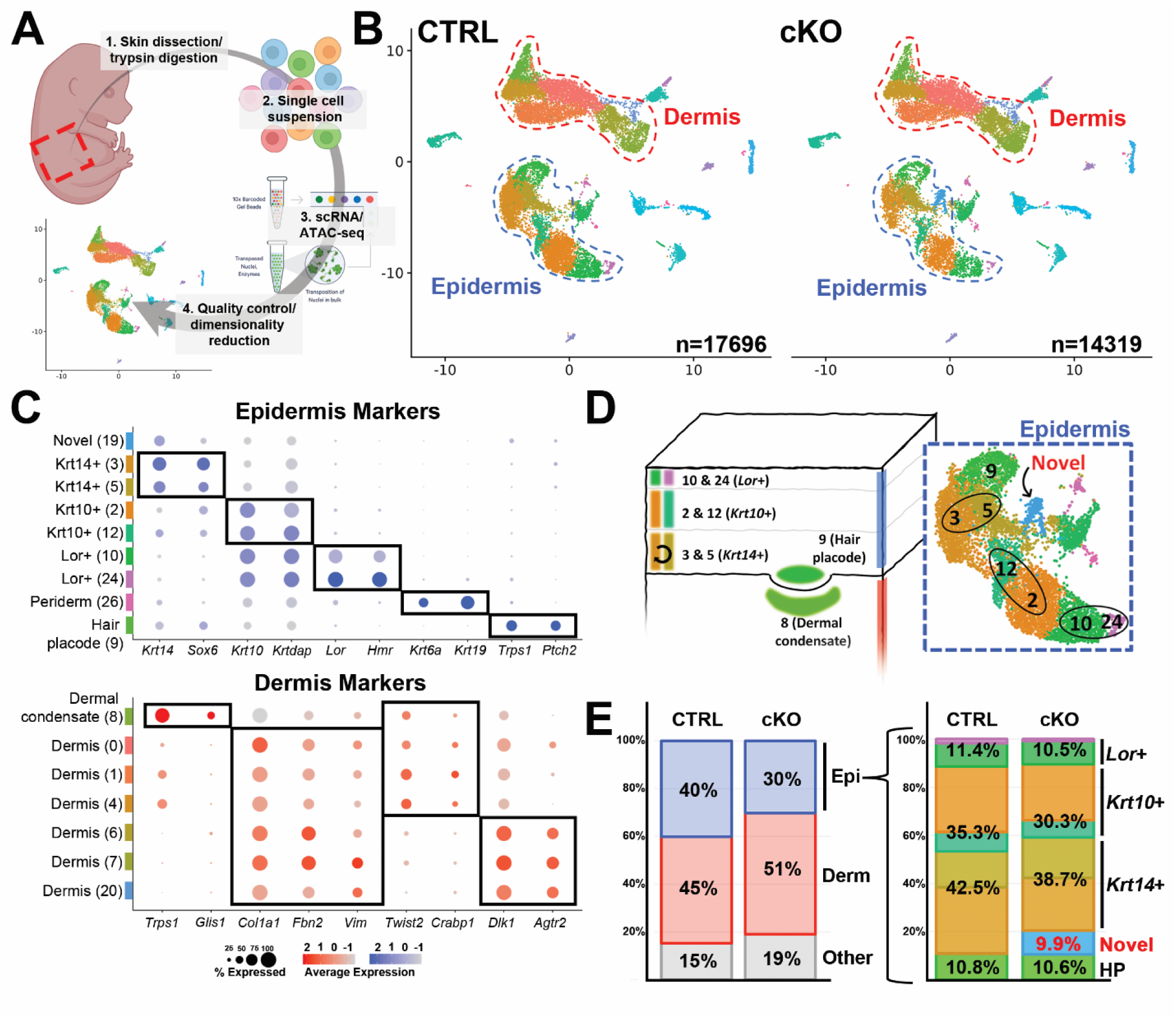
Single cell profiling of E15.5 *Prmt5* deficient epidermis reveals epidermal cell reduction coupled with emergence of a novel cell type. (**A**) A simplified schematic highlighting the key steps involved in single cell profiling of control (CTRL) and *Prmt5* cKO skin. (**B**) Unsupervised WNN clustering and UMAP projection of all cells collected from control (CTRL, 17,696 cells) and *Prmt5* cKO (cKO, 14,319 cells) samples. Note, although clustering was performed on all 32,015 cells combined, UMAPs are split by genotype. The two major subgroups based on gene expression profiles, dermis, and epidermis, are indicated. (**C**) Dot plots highlighting genes known to mark distinct subgroups, based on previous studies, of the epidermis (top, blue) and dermis (bottom, red). (**D**) Schematic and isolated UMAP (inset) highlighting the major epidermal clusters, based on gene expression profiles, along with the ‘novel’ cell population only present in *Prmt5* cKO samples. Note, the UMAP includes an overlay of both CTRL and *Prmt5* cKO cells. (**E**) Bar charts displaying the percent makeup of cell types in CTRL and *Prmt5 cKO* datasets. On the left, clusters are divided into 3 major groups, epidermis (blue), dermis (red), and other (grey), with the percentage of each indicated. On the right, subpopulations of the epidermis—divided into *Krt14*+, *Krt10*+, *Lor*+, hair placode (HP), and ‘novel’—are shown as a percentage of the total epidermal population.

Using Seurat and Signac dimensionality reduction tools (Hao et al., 2021, Stuart et al., 2021), we parsed our cell population into 30 distinct clusters (Fig. 4B). Most of these cells fell into either the epidermal (Fig. 4B, blue dashed lines, Supp. Fig. 5) or dermal (Fig. 4B, red dashed lines, Supp. Fig. 6) lineage, with other accessory cell types including things such as melanocytes, blood, and immune cells (Fig. 4B, Supp. Fig. 7). Using well characterized markers, we sorted the epidermal lineage into *Krt14*+ basal (Clusters 3 & 5), *Krt10*+ spinous (Clusters 2 & 12), and *Lor*+ differentiated (Clusters 10 & 24) keratinocyte populations (Fig. 4C, D). Additional epidermal-derived lineages such as the *Ptch2*+ developing hair follicle placode (Cluster 9) and the remnants of the *Krt19*+ periderm (Cluster 26) were also identified (Fig. 4C, Supp. Fig. 5). Unexpectedly, an additional cluster (Cluster 19) that grouped with the epidermis, was present only in *Prmt5* cKO samples (Fig. 4B, D, discussed in Fig. 5). While dermal markers are less well characterized, using recent datasets (Jacob et al., 2023), we were nonetheless able to separate the dermal lineage into three broad categories: *Twist2*+ upper dermis (Clusters 0, 1, 4), *Dlk1*+ lower dermis (Clusters 6, 7, 20), and the *Glis1*+ developing dermal condensate (Cluster 8) (Fig. 4C, Supp. Fig. 6).

**Figure 5:**
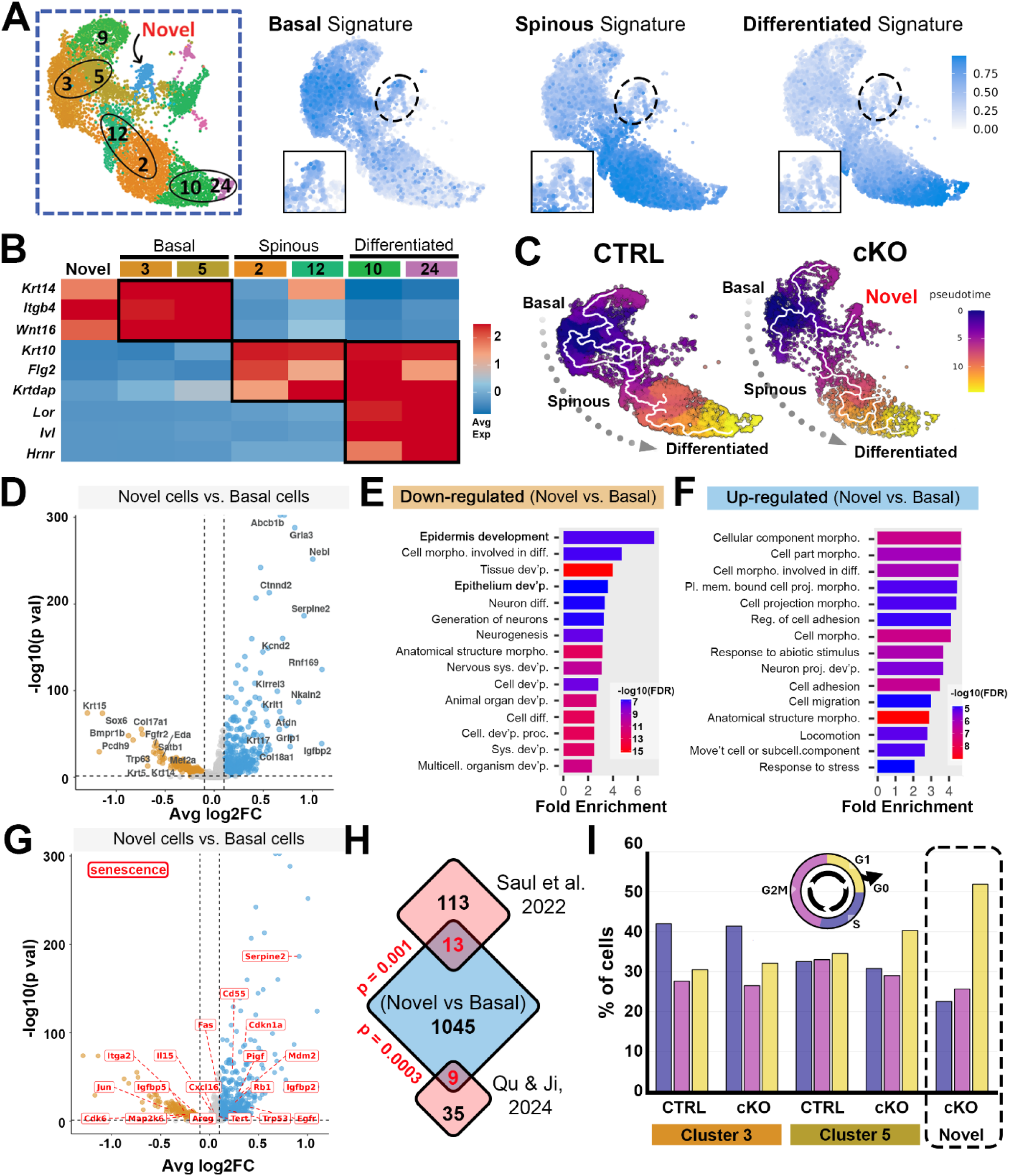
Novel epidermal cells in *Prmt5* cKOs resemble atypical basal keratinocytes and display a senescent-like profile. (**A**) UCell generated gene signature plots highlighting gene expression overlap of the top 20 expressed genes from control (CTRL) basal (Cluster 3 & 5), spinous (Cluster 2 & 12), and differentiated (Cluster 10 & 24) clusters overlaid on the epidermis focused UMAP. Intensity of blue shading represents the cumulative expression of the 20 gene signature. The inset near each ‘signature’ highlights the novel cluster (circled in the full UMAP) in higher detail. (**B**) Heatmap of select genes (rows) and their average expression level (red = high, blue = low) in distinct epidermal clusters (rows). (**C**) Monocle-derived pseudotime trajectory plots of epidermal clusters in CTRL and *Prmt5* cKO UMAPs. Note, the *Prmt5* cKO UMAP has been slightly reduced in size to better mirror the CTRL UMAP. (**D**) Volcano plot highlighting genes that are significantly (p-adj < 0.05, log2 fold change > 0.1 or < −0.1) up- (blue) or down- (gold) regulated in *Prmt5* cKO ‘novel cells’ relative to all basal cells (i.e., cluster 3 and 5 in CTRL and cKO samples). (**E, F**) GO-term enrichment analysis of the genes significantly down- (E) or up- (F) regulated in *Prmt5* cKO novel cells relative to all basal cells. (**G**) Volcano plot, as in D, but highlighting senescent associated genes with an adj p value < 0.05 in *Prmt5* cKO novel cells relative to all basal cells. (**H**) Venn diagram highlighting the number of ‘novel vs all-basal-cell’ differentially expressed genes (p_adj < 0.05, n = 1,045 total) overlapping with two independent senescent-associated gene lists (Saul et al., 2022 and Qu & Ji, 2024). p-values based on a hypergeometric test. (**I**) Histogram displaying the percentage of basal (Cluster 3 & 5) and novel (Cluster 19) cells in different phases of the cell cycle.

Having identified the major cell types in both CTRL and *Prmt5* cKO datasets (expression profiles summarized in Supp. Table 1), we next assessed the relative abundance of each cell type between genotypes. First, focusing on broad groups, we identified a reduction in the abundance of epidermal cells, relative to all cell types (i.e., epidermal, dermal and other) in the *Prmt5* cKO dataset compared to the CTRL dataset (30% vs 40%, respectively, equaling a 25% relative reduction) (Fig. 4E, left, blue). This reduction was accompanied by a corresponding increase in the relative abundance of dermal (51% vs 45%, red) and ‘other’ (19% vs 15%, grey) cell types in the *Prmt5* cKO dataset (Fig. 4E, left). Thus, although not apparent by histology (Fig. 3A, B), single cell sequencing indicates *Prmt5* cKOs at E15.5 have a slight reduction in the total number of epidermal cells relative to CTRLs. Second, given conditional deletion of *Prmt5* within the epidermis, we next assessed the relative distribution of cells among epidermal subtypes. Here we found that all major keratinocyte populations were slightly reduced in *Prmt5* cKOs relative to in CTRLs (∼4% in *Krt14*+ basal keratinocytes, ∼5% in *Krt10*+ spinous keratinocytes, and ∼1% in *Lor*+ differentiated keratinocytes) (Fig. 4E, right). Interestingly, this ∼10% relative reduction in keratinocytes from various subgroups was completely accounted for by the presence of about ∼10% of epidermal cells in the novel cluster unique to the *Prmt5* cKOs.

### While few genes are differentially expressed between canonical clusters in *Prmt5* cKO and CTRLs, a distinct novel cluster of atypical basal keratinocytes is present in *Prmt5* cKOs

Given PRMT5’s reported role in gene regulation (Stopa et al., 2015)—and the severe skin defects associated with its loss (Fig. 2)—we evaluated differentially expressed genes (DEGs) between CTRL and *Prmt5* cKO in each cluster separately. Surprisingly, within the epidermal clusters present in both genotypes, the number of DEGs was quite small (Supp. Table 2). However, this small subset of genes, including upregulation of *Dlk1* and *Rps2* and downregulation of *Ly6d* and *Rps12*, was consistently dysregulated between canonical keratinocyte populations, in *Prmt5* cKOs versus CTRLs (Supp. Fig. 8, Supp. Table 2). Correspondingly, minimal GO-terms were enriched among down or upregulated DEGs, when grouped by basal (Clusters 3 & 5), spinous (Clusters 2 & 12), or differentiated (Clusters 10 & 24) clusters (Supp. Fig. 8). It is interesting to note that multiple genes were found to be upregulated in dermal clusters of *Prmt5* cKO’s, relative to CTRLs, presumably due to cell non-autonomous effects (Supp. Fig. 9, Supp. Table 2). Few gene expression changes were detected in other clusters, consistent with unchanged levels of PRMT5 in them (Supp. Fig. 10, Supp. Table 2). Correspondingly, almost no significant changes (p_adj < 0.05) in chromatin accessibility were observed between *Prmt5* cKO and CTRL clusters (Supp. Fig. 11, Supp. Table 3). These results indicate that, within clusters present in both genotypes, minor changes in gene expression are unlikely to be mediated by changes in chromatin accessibility.

We next turned our attention to the novel cluster only present in the *Prmt5* cKO dataset (i.e., Cluster 19). Based on this cluster’s location in UMAP space (i.e., proximity to other clusters), we hypothesized that its molecular profile was most like that of basal keratinocytes (Fig. 5A). To explore this possibility, we used the UCell package (Andreatta and Carmona, 2021) to plot a basal, spinous, and differentiated ‘gene signature’ (i.e., the top 20 genes representing each group) on the epidermal UMAP. Consistent with our hypothesis, the ‘basal gene signature’ appeared to overlap most with the novel cluster, relative to the spinous and differentiated ‘signature’ (Fig. 5A). Gene expression analysis illustrated the overlap of basal keratinocyte genes (e.g., *Krt14*, *Itgb4*, *Wnt16*) and the exclusion of common spinous (e.g., *Krt10*, *Flg2*, *Krtdap*) and differentiated (e.g., *Lor*, *Ivl*, *Hrnr*) genes within the novel cluster (Fig. 5B). Pseudotime analysis of epidermal clusters revealed the expected progression from basal to spinous to differentiated keratinocyte clusters in both CTRL and *Prmt5* cKO datasets. However, within the *Prmt5* cKO dataset, there was an abnormal trajectory from the normal basal keratinocyte cluster to the novel keratinocyte cluster, instead of to the spinous lineage (Fig. 5C). In sum, these data suggest that the novel cluster is an abnormal derivative of basal keratinocytes.

We next examined what molecular features made cells in the novel cluster distinct from basal keratinocytes. We assessed the DEGs between the novel cluster and the basal keratinocyte clusters in both CTRL and *Prmt5* cKO profiles (which had very few DEGs between them) (i.e., ‘All Basal’). This analysis revealed 180 down- and 865 up-regulated DEGs in the novel cluster relative to the normal basal cluster (Fig. 5D, Supp. Table 3). Among the down regulated genes were several known to regulate basal keratinocytes (e.g., *Krt14*, *Col17a1*, *Eda*, *Sox6*, *Satb1*), and consistent with this observation, the down-regulated genes were enriched for gene ontology (GO) terms ‘Epidermis Development’, ‘Tissue Development’, and ‘Epithelium Development.’ Genes whose expression was up regulated in the *Prmt5* cKO-associated novel cells relative to in basal keratinocytes (e.g., *Igfbp2*, *Serpine2*, *Rnf169*, *Abcb1b*, *Krt17*) were enriched for the GO terms ‘Cell Morphology’, ‘Cell Projection’, ‘Cell Adhesion/Migration’ and ‘Response to Stress’. Independent bulk RNA-sequencing of skin from E15.5 CTRL and *Prmt5* cKO embryos confirmed several of these up- and down-regulated genes (of 649 overlapping significant DEGs, 71% changed in the same direction) (Supp. Fig. 12).

### The *Prmt5* cKO-associated novel cluster is enriched for a *Cdkn1a*-driven senescence phenotype, including expression of SASP molecules and stress keratins

Although a defined list of senescence associated genes is often context dependent and thus omitted from GO term enrichment analysis, several recent studies have proposed a core list of senescence markers (Qu et al., 2024, Saul et al., 2022). Evident from our differential expression analysis between novel and basal cell clusters, was a significant overrepresentation of these genes (e.g., *Cdkn1a/p21*, *Igfbp2*, *Serpine2*, *Trp53*, *Rb1*, etc.) (Fig. 5G). For example, of a 126 senescence-associated gene list (Saul et al., 2022), 13 genes overlapped our ‘novel vs basal’ DEG list (hypergeometric test, p = 0.001) and of a separate 44 senescence-associated gene list (Qu et al., 2024), 9 genes overlapped (hypergeometric test, p = 0.0003) (Fig. 5H). Additionally, cells in the *Prmt5* cKO-associated novel cluster showed a decreased propensity for being in active phases of the cell cycle (S or G2M) relative to cells in the basal cell clusters (Fig. 5I). Together these results indicate that the *Prmt5* cKO-associated novel cluster is comprised of basal keratinocytes with elevated levels of senescence.

Two hallmarks of cell senescence are expression and activation of *Cdkn1a* (*p21*)—a cyclin-dependent kinase inhibitor that tightly regulates cell cycle progression (Yan et al., 2024)—and execution of a ‘senescent associated secretory phenotype’(SASP) (Xu et al., 2024). Within the *Prmt5* cKO novel cluster, expression of *Cdkn1a* was modestly but significantly elevated relative to in basal keratinocytes (FC = 1.28, p_adj <0.0001) (Fig. 6A). This increase was reflected in more intense anti-CDKN1A immunofluorescent signal in the epidermis of *Prmt5* cKOs compared to CTRLs (Fig. 6B, C). Interestingly, scATAC-seq data revealed a trend towards increased chromatin accessibility at the *Cdkn1a* promoter in the novel cluster relative to in basal keratinocytes, although this increase did not reach significance (FC= 1.79, p_adj =0.51) (Fig. 6D, Supp. Table 3). Similarly, *Igfbp2*—encoding a secreted protein and component of the SASP response (Evans et al., 2024)— was expressed at higher levels in cells of the novel cluster relative to basal cells (Fig. 6E) (FC = 2.13, p adj < 0.0001) and again this was reflected in more intense anti-IGFBP2 immunofluorescent signal at E15.5 in both epidermis and dermis of *Prmt5* cKO embryos compared to CTRLs (Fig. 6F, G). Finally, among other stress related genes (Fig. 5D, Supp. Fig. 13), expression of the stress-induced keratin gene, *Krt17*, was elevated in the novel cluster relative to in basal cells (FC = 1.46, p adj < 0.0001) (Fig. 6H). Immunofluorescent analysis revealed expanded expression of KRT17 throughout *Prmt5* cKO basal keratinocytes, relative to KRT17’s more restricted expression in CTRLs, particularly at later stages of phenotype progression (Fig. 6I, J).

**Figure 6:**
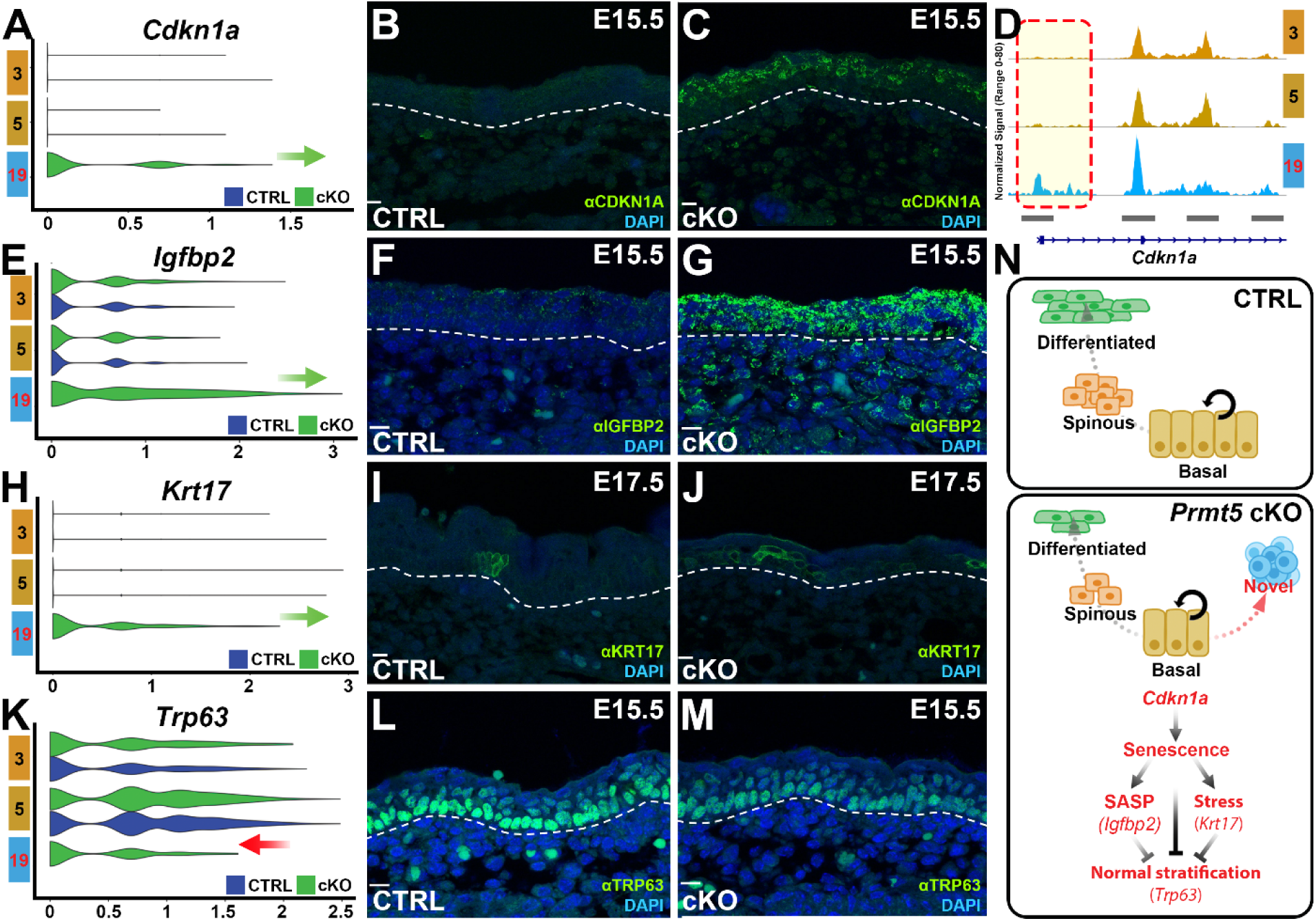
The *Prmt5* cKO novel cluster represents a *Cdkn1a*-driven senescent population which is confirmed *in vivo*. (**A-M**) scRNA-seq violin plots for *Cdkn1a* (A), *Igfbp2* (E), *Trp63* (H), and *Krt17* (K) in the novel (19) and basal (3/5) keratinocyte clusters. Corresponding α-CDKN1A (B-C), α-IGFBP2 (F-G), α-TRP63 (I-J), and α-KRT17 (L-M) immunofluorescence from E15.5 (B, C, F, G, I, J) or E17.5 (L, M) CTRL (B, F, I, L) or *Prmt5* cKO (C, G, J, M) embryos. Integrative Genomics Viewer (IGV) reveals a suggestive (FC=1.74, p_ adj=0.51) locus specific increase in chromatin accessibility at the *Cdkn1a* promoter (D). (**N**) A schematic summarizing the normal progression of keratinocytes in CTRL (top) and atypical progression in Prmt5 cKOs (bottom), including the molecular features involved. Sections shown are the craniofacial epidermis, lateral to the oral cavity, in a coronal plane. White dashed lines demarcate the boundary of the dermis and epidermis. Nuclei are counterstained with DAPI. All section immunofluorescence images are Z-stacks compiled with ‘Max Intensity’ in FIJI. Scale bar = 10 µm.

We also examined expression and distribution of anti-TRP63 immunoreactivity, a master transcriptional regulator of the keratinocyte fate (Yang et al., 1999). *Trp63* expression was lower in cells within the novel cluster relative to in basal cells (FC = 0.65, p_adj <0.0001) (Fig. 6K), which was reflected in less intense anti-TRP63 immunofluorescent signal in the basal cells of *Prmt5* cKOs, relative to CTRLs (Fig. 6L, M). Interestingly, a subset of basal keratinocytes with a more pronounced reduction in TRP63 expression was apparent, potentially representing cells defined by the novel cluster (Fig. 6L, M). Collectively, these findings correlate loss of *Prmt5* in basal keratinocytes with elevated senescence, increased cell stress, and reduced TRP63.

In summary, we propose a model in which loss of *Prmt5* in the epidermis drives an increase in *Cdkn1a*—potentially by relieving epigenetic repression—causing a subset of basal keratinocytes to lose their proper developmental trajectory towards spinous keratinocytes and instead gain an abnormal senescence phenotype (Fig. 6N). This senescence triggers the secretion of SASP factors, particularly IGFBP2, and upregulates stress keratin expression, including *Krt17*. Collectively, these changes disrupt proper epidermal stratification, as evidenced by a decrease in *Trp63* expression and corresponding skin phenotypes in *Prmt5* cKO embryos.

## DISCUSSION

Here, we showed that PRMT5 is expressed throughout the developing skin and that introduction of the CRE recombinase encoding *Crect* allele, in conjunction with the *Prmt5* flox allele, causes a profound reduction in *Prmt5* mRNA and protein levels solely within the epidermis at early (E13.5, E15.5) stages. However, we did observe a mild reduction in late-stage (E17.5) PRMT5 dermal expression, where CRE does not appear to be expressed. We also found that skin of *Prmt5* cKO embryos appears histologically normal until approximately E15.5, when the epidermis exhibits increasing hyperplasia. By E18.5, *Prmt5* cKOs had a reduced basal layer, lacked spinous and granular layers, and had a sparse cornified layer, relative to CTRL. We conducted single cell multiome analysis of *Prmt5* cKOs and CTRLs at E15.5 when the histological features are first starting to emerge. Analysis of typical clusters (e.g., basal, spinous, differentiated) detected few differentially expressed genes (DEGs) and few differentially accessible chromatin loci (DACs), between genotypes. However, this analysis revealed a subset of basal cells within *Prmt5* cKOs enter an atypical state characterized by expression of markers of senescence and cell stress. Below we focus on several implications of these findings.

First, cellular and molecular profiling of *Prmt5* cKOs—at the apparent onset of epithelial phenotypes—revealed differences in cell identity and gene expression. Overall, we observed a modest decrease in total keratinocyte populations, with a reduction coming from each canonical (e.g., basal, spinous, differentiated) layer. This reduction coincided with the emergence of a novel or atypical cell population that closely clustered with other keratinocytes. The novel cell population exhibited a gene signature most similar to the stem cell-like basal population, including presence of canonical basal markers, such as *Krt14* and *Itgb4,* and absence of more superficial markers, such as *Krt10*, *Flg2*, *Lor*, and *Ivl*. Additionally, developmental trajectory analysis revealed that this population likely diverged directly from the basal cells. Fittingly, these cells exhibited a more divergent gene expression profile than any other CTRL to *Prmt5* cKO cluster comparison. Consistent with PRMT5’s known role in transcriptional repression (Karkhanis et al., 2011), the majority of DEGs associated with the atypical cluster were significantly upregulated (∼83% of the 1,045 DEGs). Included within this group were genes associated with cell morphology, adhesion, and senescence. The latter was significantly over-represented and consistent with this, cells in the atypical cluster had a propensity to not be actively cycling, relative to basal cells. Thus, the absence of PRMT5 in the ectoderm resulted in progression of basal keratinocytes down an atypical path, creating a novel cluster marked by senescence.

Equally intriguing was that despite the presence of this novel cluster, a substantial number of cells progressed down a ‘normal’ developmental trajectory, at least until E15.5, in *Prmt5* cKO skin. Differential gene expression between CTRL and *Prmt5* cKOs revealed few statistically significant dysregulated genes in these canonical clusters. Given the presence of these ‘normal’ keratinocytes and their similarity to respective CTRLs, it is possible a compensatory mechanism enables some keratinocytes to retain normal expression profiles and developmental trajectories under *Prmt5* cKO conditions. Given the presence of multiple PRMT proteins in keratinocytes (Bao et al., 2017), it is plausible that another *Prmt* is upregulated upon loss of *Prmt5* to preserve sufficient levels of symmetric methylation. A non-mutually exclusive possibility is that PRMT5 function is not necessary until ∼E15.5 (or just prior), and thus a first phase of stratification has occurred that is PRMT5-independent. Two phases of basal keratinocyte maturation have been proposed, with the timing coinciding with the onset of the *Prmt5* cKO pathology (Damen et al., 2021). Under such a scenario, we predict that the novel cluster would continue to increase at the expense of the canonical clusters as embryonic development proceeds (e.g., if conducting single cell analysis at later embryonic timepoints). Further, targeting *Prmt5* deletion in basal keratinocytes at a stage later than *Crect*, such as with *K14*:CRE (Andl et al., 2004, Dassule et al., 2000), should result in a similar skin phenotype.

Second, the cellular mechanism explaining progressive loss of suprabasal keratinocytes after E15.5 in *Prmt5* cKOs is unclear. Evidence from cultured keratinocytes indicates that PRMT5 suppresses keratinocyte differentiation stimulated by PKCδ and p38δ-mediated signaling (Kanade and Eckert, 2012). We therefore anticipated premature differentiation of keratinocytes in *Prmt5* cKOs, which could potentially result in thinner than normal skin. However, analysis of canonical markers of each epidermal layer (KRT14, KRT10, and IVL) revealed grossly correct timing, expression, and localization. In addition, levels of apoptosis and proliferation were comparable in epidermal cells of *Prmt5* cKO and CTRLs. Given that the skin is an external organ in continuous contact with the amniotic fluid in utero, we cannot rule out that in the skin of *Prmt5* cKO after E15.5 a portion of keratinocytes, perhaps those derived from the novel cluster—expressing signs of senescence and stress—lose adhesion (Kato et al., 2021) (Supp. Fig. 13) and are therefore shed into the environment. It will be critical to apply lineage labeling techniques in *Prmt5* cKOs to track progression of basal—and subsequent suprabasal— cells during epithelial development.

Third, the relevant substrate of PRMT5 mediated methylation underlying changes in mutant skin remains unclear. Known PRMT5 substrates that regulate keratinocyte differentiation include the transcription factor KLF4 (Hu et al., 2015), an unknown component of a complex that includes p38δ and PKCδ (Kanade and Eckert, 2012), and the histones H3 and H4 (Saha et al., 2016). As changes in histone methylation may affect chromatin accessibility, we performed scRNA and scATAC-seq analyses on bulk skin extracts from E15.5 embryos, just prior to phenotype development. Comparing the scATAC-seq profiles in canonical (e.g., basal, spinous, differentiated) clusters between CTRL and *Prmt5* cKO revealed very few statistically significant changes in chromatin accessibility (Supp Fig. 11). However, comparing these profiles between the atypical basal cell cluster in *Prmt5* cKO to the typical basal clusters in either *Prmt5* cKO or CTRLs, revealed ∼60 significantly more accessible ATAC peaks, with additional peaks, including the *Cdkn1a* promoter, approaching significance (Supp. Table 3, Supp Fig. 11, Fig 6D). It is important to note that the sequencing depth in scATAC-seq may only be sufficient to detect large differences in ATAC signal. Currently, it is still unclear whether these PRMT5-dependent changes in chromatin accessibility result from altered methylation of histones, nor whether the minimal changes contribute to the overt defects in skin development. Beyond roles in the nucleus, PRMT5 is known to also localize to the cytoplasm under different developmental and oncogenic contexts (Gu et al., 2012, Wang et al., 2015). Given that PRMT5 has a trend towards cytoplasmic enrichment at E15.5, PRMT5’s methyltransferase function may act primarily on cytoplasmic proteins, at this stage. Deeper sequencing, perhaps in bulk ATAC-seq of keratinocytes, and evaluation of methylation of various PRMT5 substrates in *Prmt5* cKOs and CTRLs will help to resolve the relevant PRMT5 substrate in this context.

Finally, in our proposed model (Fig 6N), we hypothesize that PRMT5 directly represses *Cdkn1a* expression, potentially through epigenetic mechanisms, leading to *Cdkn1a* overexpression and a subsequent increase in cell senescence in *Prmt5* cKOs. In turn, SASP-components, such as IGFBP2, can stabilize CDKN1A in keratinocytes, potentially creating a positive feedback loop that amplifies the senescence pathway (Mercurio et al., 2020). However, it remains possible that an intermediate factor is responsible for the link between *Prmt5* loss and aberrant *Cdkn1a* expression. Cellular senescence can be triggered by various environmental stressors, including DNA damage, reactive oxygen species production, and endoplasmic reticulum stress (Huang et al., 2022). Notably, PRMT5 is involved in the DNA damage response (Yin et al., 2023) and proper RNA splicing (Muro et al., 2024), and both processes have key marker genes upregulated in the novel cluster (i.e., DNA damage-associated *Rnf169* & *Ercc5,* and RNA splicing-associated *Hnrnph1* & *Sf3b2*) (Supp. Fig. 13). Therefore, it is plausible that *Prmt5* loss results in increased DNA damage or splicing errors, inducing cellular stress, and triggering senescence indirectly. Further studies, including DNA damage profiling, alternative splicing analysis, protein interaction assays, and rescue experiments, are needed to determine whether PRMT5 regulates *Cdkn1a* directly or indirectly.

In summary, our findings suggest that PRMT5 is essential for preventing senescence in basal keratinocytes, with its absence leading to improper stratification, defective barrier formation, and perinatal lethality. These results provide insights into the epigenetic regulation of basal keratinocyte maintenance and senescence and may inform future studies on the role of epigenetics and senescence in congenital skin disorders.

## ACKNOWLEDGEMENTS

We acknowledge the University of Iowa Central Microscopy Research Facility and Genomics Division for their technical support with tissue processing and scMultiome-seq experiments, respectively. These facilities are funded through user fees and generous financial support from the Carver College of Medicine. We also thank Dr. Martine Dunnwald from the Department of Anatomy and Cell Biology for her invaluable expertise and guidance in skin development; Dr. Shareef Dabdoub from the College of Dentistry Division of Biostatistics and Computational Biology, Dr. Colin Kenny from the Department of Surgery, Dr. Nate Mullin, and Jerzy Twarowski for their insights into coding and scMultiome-seq analyses; and Dr. Bin He, Department of Biology, Andrew Frank, Department of Anatomy and Cell Biology, members of the Van Otterloo lab, as well as our colleagues in the Craniofacial Interest Group, for their invaluable feedback on this study. Finally, we thank Dr. Steven Vokes at the University of Texas, Austin and Dr. Trevor Williams at the University of Colorado, Anschutz Medical Campus, for providing generous access to the *Prmt5* conditional and *Crect* allele, respectively.

## MATERIALS & METHODS

### Mouse Strains and Crosses

Mouse experiments were approved by the Institutional Animal Care and Use Committee at The University of Iowa (Protocol #2032197). A *Prmt5* conditional mouse line *Prmt5*^tm2c(EUCOMM)Wtsi^ (referred to as *Prmt5*^c^, Norrie et al. 2016) was crossed with *Crect*^+/−^ males (Schock et al. 2017) to generate *Crect*^+/−^;*Prmt5*^c/+^ males. These were crossed with *Prmt5*^c/c^ females to generate *Crect*^+/−^;*Prmt*5^c/c^ embryos (*Prmt5* cKOs). *Prmt5* alleles were detected with primers 5’- TTCTGGCCTCCATGGGGGAA -3’ and 5’- TGGAACTGCAGGCATATGCC -3’, generating a 365bp fragment for the conditional allele and a 216bp fragment for the wild-type allele. *Crect* alleles were detected with primers 5’- CCTCACTGATCCACATAT GTCCTTCCGAAAGCTGC -3’, 5’- GATGCTAGAAAGCTGAGGCTGGGCTTAGCTTGCTAGGC-3’ and 5’- CTACGCCGCGAACTTGCTTCTAGAGCG -3’, generating a 385bp fragment for the conditional allele and a 586 bp for the wild-type allele. *Crect*^+/−^ mice were crossed with the *mTmG*^+/+^ (Muzumdar et al. 2007) line or with the *R26R*^+/+^ (Soriano 1999) line and used to validate Cre expression via GFP or β-galactosidase, respectively.

### Toluidine Blue Permeability Assay

Embryos were dissected, washed in 100% methanol, and rehydrated in PBS. Embryos were then incubated in 0.1% Toluidine Blue (Millipore Sigma, T3260) in PBS for 2 hours at 4°C. Stain was washed off with successive PBS incubations prior to imaging. White balancing was performed using GIMP.

### Skeletal Staining

Dissected embryos were stripped of excess skin, viscera, and adipose tissue then incubated in 95% ethanol for 3 days and 100% acetone for 2 days to further remove tissue. Next, embryos were incubated in staining solution (0.03% alcian blue 8 GX, 0.1% alizarin red S, 5% glacial acetic acid, 70% ethanol) for 3 days at 37°C. De-staining was performed via incubation in 2% KOH for 1-2 days at room temperature. Embryos were transferred to glycerol for storage and imaging.

### Tissue embedding and sectioning

Embryos were dissected, beheaded, and fixed overnight at room temperature in 4% PFA. For paraffin sections, samples were dehydrated in a gradient of ethanol (e.g., 70%, 90%, 95%, 100%) and then xylene, before being embedded into paraffin wax. Embryos were then sectioned at 7 µm on a manual microtome (Thermofisher, HM 325) before being fixed to slides. For frozen sections (α-GFP and α-CDKN1A), fixed samples were embedded in optimum cutting temperature (O.C.T.) compound (Tissue-Tek) and 10 µm sections were prepared using a Leica CM3050X Cryostat.

### Hematoxylin & Eosin Staining

For histology, paraffin sections were incubated for 3 minutes in hematoxylin and 30 seconds in eosin before being de-stained via acid ethanol. Coverslips were adhered using Permount (Fisher Scientific, SP15-100) prior to imaging.

### Immunofluorescence

Sections were incubated in primary antibody overnight at 4°C. Primary antibodies include: α-CDKN1A (1:50, ProteinTech, 10355-1-AP), α-CLDN1 (1:60, Abcam, ab15098), α-COL4 (1:500, Invitrogen, PA1-85320), α-eGFP (1:500, Novus Biologicals, NB600-308), α-FLG (1:500, Biolegend, 905804), α-IGFBP2 (1:200, Bioss, bs-1108R), α-KI67 (1:200, Abcam, ab16667), α-KRT6 (1:500, Biolegend, PRB-169F), α-KRT10 (1:500, Biolegend, 905404), α-KRT14 (1:20, SantaCruz, sc-53253), α-KRT17 (1:1000, Proteintech, 17516-1-AP), α-NF (0.5ug/ml, Hybridoma, AB531793), α-PRMT5 (1:200, Millipore, 07-405), α-TRP63 (1:500, Cell Signaling, 13109S), and α-VCL (1:300, Sigma, V9131). Sections were washed and then incubated with the corresponding alexafluor-conjugated secondary antibody for ∼1 hour, at room temperature. Slides were counterstained with 300nM DAPI stain solution (ThermoFisher, D1306) for 5 min at room temperature and coverslips were adhered with ProLong Mountant with DAPI (Invitrogen, P36941). α-PRMT5 antibody-labeled sections were also processed using the Vector TrueVIEW Autofluorescence Quenching Kit (Vector Labs, SP-8400-15) following manufacturer’s instructions.

### TUNEL staining

TUNEL staining was performed using the DeadEnd Fluorometric TUNEL System (Promega) following manufacturer’s instructions.

### Slide Imaging

Slides were imaged on either a Zeiss 700 or Zeiss 880 confocal microscope using either 20X or 40X magnification. Z-stack images were collected through the sample and were processed using ZEN software and ImageJ.

### Western Blot

E18.5 skin was dissected from embryos and incubated with 1mg/ml Dispase II (ThermoFisher) in Hepes Buffer (20 mM Hepes, 115 mM NaCl, 1.2 mM CaCl2, 1.2 mM MgCl2, 2.4 mM K2HPO4) for 30 min at 37°C, prior to manual separation of the epidermis and dermis. Samples were homogenized in RIPA buffer (50mM Tris pH 7.4, 150mM NaCl, 0.1% SDS, 0.5% sodium deoxycholate, 1% NP-40) containing complete mini protease inhibitor (Roche). Approximately 5-10µg of lysate was ran on 4-12% NuPAGE Bis-Tris Gels (ThermoFisher), transferred to PVDF membranes, and incubated overnight at 4°C with the primary antibody. Primary antibodies included: α-PRMT5 (1:10,000, Abcam, ab109451) and α-GAPDH (1:1000, SantaCruz, sc-390257). Membranes were then incubated with the corresponding goat-anti-rabbit (1:20,000, Licor IR Dye 800CW Anti-Rabbit, 926-32211) or goat-anti-mouse (1:20,000, Licor IR Dye 800CW Anti-Mouse, 926-32212) secondary antibody for 1 hour at room temperature and imaged using the Licor Odyssey M Imager.

### Single cell suspension and ATAC reaction

For single cell multiome-sequencing experiments, embryos were collected at E15.5 from multiple litters. Yolk-sac tissue was lysed quickly using the Extract-N-Amp Tissue PCR Kit (Sigma-Aldrich) and embryos quickly genotyped using Phusion High-Fidelity DNA Polymerase (New England Biolabs) and shortened PCR cycling parameters. During genotyping, the back skin from all E15.5 mouse embryos was dissected and immediately transferred to ice cold DEPC-PBS (Millipore Sigma). Samples were incubated in 0.25% Trypsin (ThermoFisher) for 30-45 mins at 37°C with frequent pipette mixing. Once cells reached a single cell suspension, the reaction was quenched with 10% FBS (ThermoFisher), passed through a 40 µM cell strainer (Flomi Cell Strainers, SP Bel-Art), and triplicates of each genotype (CRE-negative controls or *Prmt5* cKOs, based on quick-genotyping results) pooled. 500,000 cells from each pooled sample were incubated with lysis buffer (10mM Tris HCl pH 7.5, 10mM NaCl, 3mM MgCl2, 0.1% NP-40) for 5 mins on ice, nuclei extracted and quantified using trypan blue.

scRNA-Seq and scATAC-Seq sequencing libraries were prepared by the University of Iowa’s Iowa Institute of Human Genetics Genomics Division Laboratory. Briefly, 10,000 nuclei were targeted for bulk transposition before individual nuclei were encapsulated in oil droplets along with the 10X GEM code beads in the Chip J cartridge (P/N: 1000234) using the 10X Genomics Single Cell iX Chromium Controller. Generation of gel beads in emulsion (GEMs), barcoding, pre-amplification PCR, and ATAC library construction were all performed as recommended by the manufacturer (10X Genomics, Chromium Next GEM Single Cell Multiome + ATAC Reagent Kit, Rev F, P/N:1000283). The gene expression libraries were pooled and sequenced on a NovaSeq 6000 to give at least 20,000 reads per nuclei. The ATAC-Seq libraries were pooled and sequenced on a NovaSeq 6000 to give at least 25,000 reads per nuclei.

### Data processing

Sequencing results were demultiplexed and converted to FASTQ format using the Illumina bcl2fastq software. scRNA and scATAC-seq reads were processed and aligned to the mm10 reference genome using the Cell Ranger Count function. Further analysis and visualization was performed using Seurat (v4.3) (Hao et al., 2021) and Signac (v1.9) (Stuart et al., 2021). Briefly, peaks were unified between conditions and filtered based on length (20-10,000). EnsDb.Mmusculus.v79 was used for gene annotations. Cells underwent manual filtering to ensure cells used were between nCount_ATAC (100-50,000), nCount_RNA (100-25,000), nucleosome_signal <4, and TSS_enrichment >1. scRNA-seq separately underwent an additional SCTransform analysis. scATAC-seq underwent FindTopFeatures, RunTFIDF, and RunSVD. Datasets were then intersected using FindIntegrationAnchors (1:30, k.anchors=5) and IntegrateData (1:30). SCTransform, RunTFIDF, FindTopReatures, and RunSVD were rerun after integration. FindMultiModal neighbors using PCA (1:50) and LSI (2:50) was performed prior to FindClusters to generate the final UMAP. Trajectory analysis was performed with Monocle3 (Trapnell et al., 2014). Gene signature scoring was performed with UCell (Andreatta and Carmona, 2021). Cell cycle analysis was performed using CellCycleScoring with the cc.genes.updated.2019 converted to mouse format by Gprofiler2 (Kolberg et al., 2020). GO ontology analysis was performed using GO Biological Process and plotted using ShinyGO 0.8 (Ge et al., 2020).

### Bulk RNA-sequencing

Back skin from E15.5 mouse embryos (3 control and 3 *Prmt5* cKO embryos) was dissected and immediately transferred to ice cold DEPC-PBS (Millipore Sigma, D5758). RNA was isolated using the RNeasy Fibrous Mini Kit (Qiagen, 74704) following manufacturer’s instructions. cDNA libraries were generated using the Illumina TruSeq Stranded mRNA kit and samples sequenced using the NovaSeq6000 platform (∼30 million, 200 bp, paired-end reads/sample). Following demultiplexing, reads were mapped to the Mm10 genome using STAR (Dobin et al., 2013), transcript and read counts conducted using Stringtie (Pertea et al., 2015), and differential gene expression analysis carried out using DeSeq2 (Love et al., 2014). This approach identified 1,454 differentially expressed genes (DEGs) (p adj < 0.01, log2FC > 1), although this list was skewed by the reduction in total ectoderm of *Prmt5* cKOs, compared to CTRLs, as bulk samples contained both epidermis and dermis. As a result of reduced epidermis tissue in the overall *Prmt5* cKO sample (relative to CTRLs), a large portion of epidermis-associated genes were correspondingly significantly down-regulated.

To compare sc-RNAseq data with bulk RNA-seq data, differentially expressed genes identified between the novel cluster and ‘all basal’ clusters (p adj < 0.05, N = 1,045 genes) were overlapped with significant DeSeq2 identified genes (p adj < 0.01, N = 5,340 genes), resulting in the 649 genes plotted in Sup. Fig. 12.

### Graphs and Statistics

Schematics were partially created using BioRender.com. Bar graphs were generated using GraphPad and XNomial was used in RStudio to assign significance. Ki67 & TUNEL quantification was performed in ImageJ by manually counting positively stained cells and dividing by the total area of Krt14 stained epidermis or the number of DAPI positive epidermal cells, respectively. Hypergeometric tests were conducted in RStudio using the *phyper* function, with overlapping of DEGs and SASP associated gene-sets conducted with a conservative estimate of 20,000 total genes in the genome.

## Abbreviations

PRMT5: Protein Arginine Methyl Transferase
cKO: conditional knockout
scRNA/ATAC-seq: single-cell RNA and ATAC sequencing
SASP: senescence associated secretory phenotype

---

**Supplementary Figure 1:**
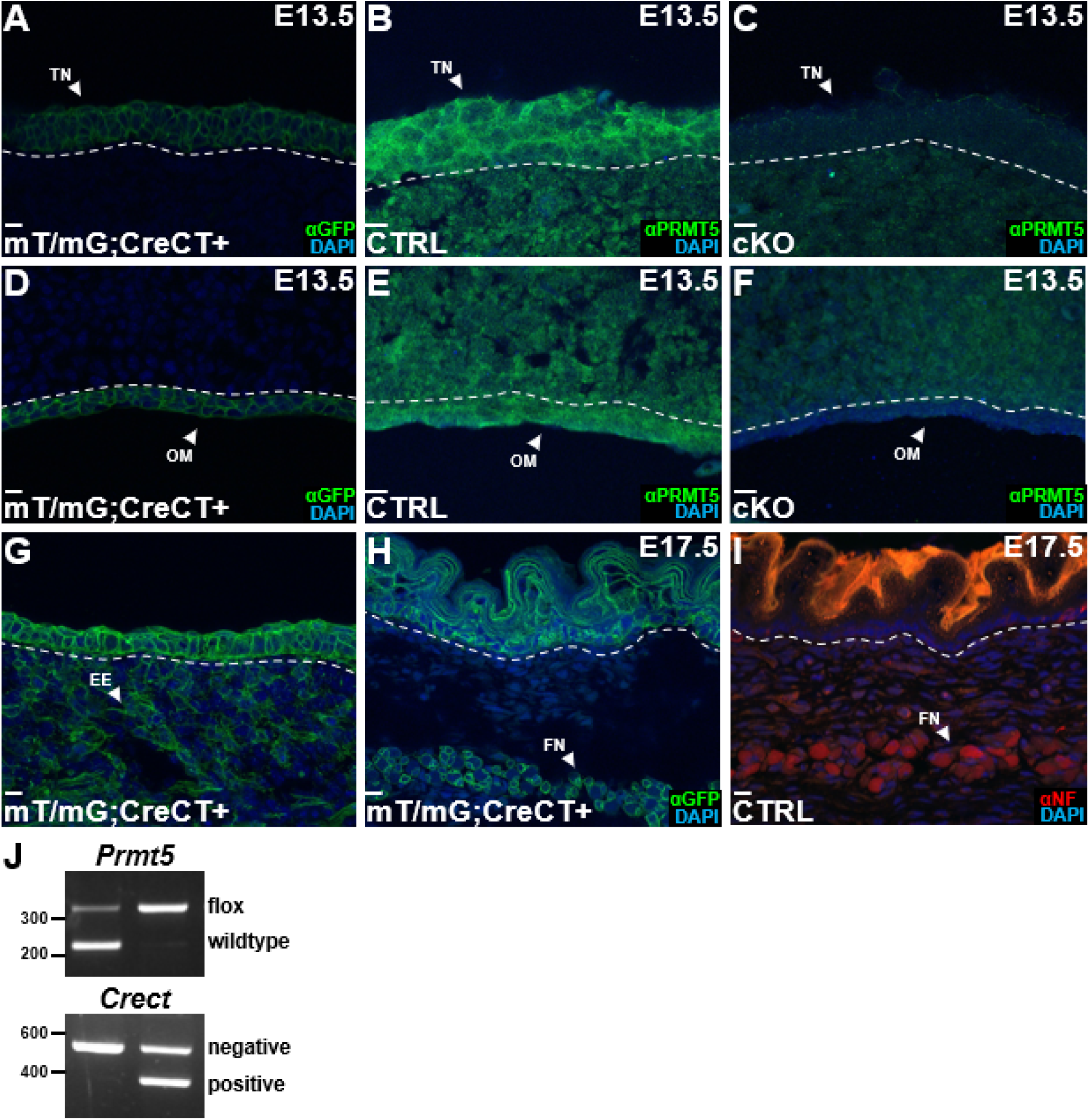
PRMT5 is expressed throughout epithelial development and can be conditionally deleted using *Crect*. (**A-I**) Coronal sections from early (E13.5) (A-F) or late (E17.5) (G-I) stage CTRL (B, E, I), cKO (C, F), or *mT/mG*;*Crect*+(A, D, G, H) embryonic heads processed by α-GFP (A, D, G, H), α-PRMT5 (B-C, E-F), or α-Neurofilament (I) immunofluorescence. (**J**) Agarose based PCR gels from tail or yolk sac samples processed using *Prmt5* or *Crect* primers (for sequences see methods). Bands shown are for *Prmt5* flox (365bp), *Prmt5* wild type (216bp), *Crect* Negative (586bp), and *Crect* Positive (385bp). Note, sections shown are lateral to the oral cavity, in a coronal plane. White dashed lines demarcate the boundary of the ectodermally-derived and mesenchyme-derived tissues. Acronyms include tongue (TN), oral mucosa (OM), ectopic expression (EE), and facial nerve (FN). Nuclei are counterstained with DAPI. All sectional immunofluorescence images are five Z-stacks compiled with Max Intensity in FIJI. Scale bar = 10µm.

**Supplementary Figure 2:**
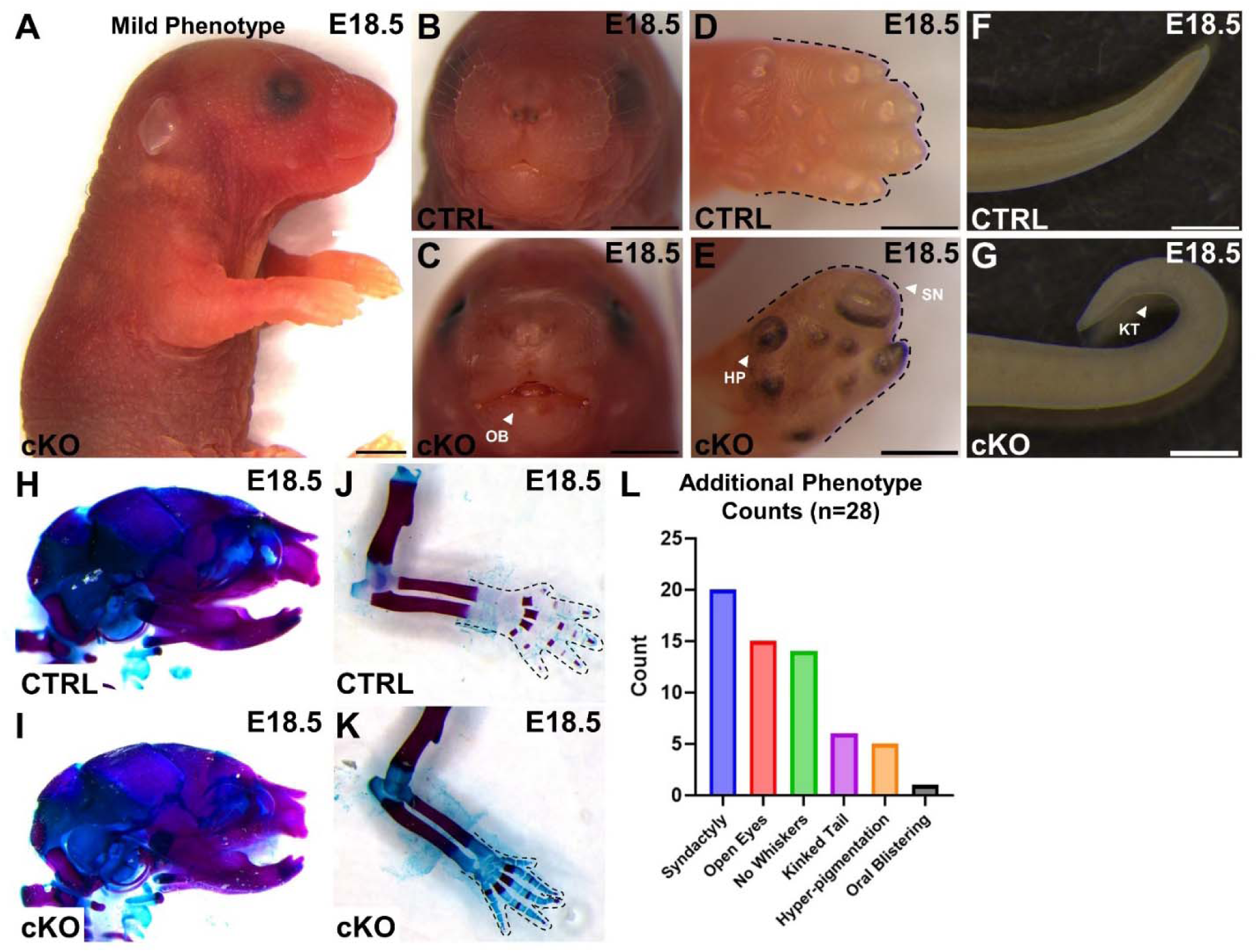
Epithelial-specific loss of PRMT5 results in a variety of epithelial-related defects but lacks obvious skeletal abnormalities. (**A-K**) Brightfield lateral (A, D-K) and frontal (B-C) view images of a CTRL (B, D, F, H, J) or *Prmt5* cKO (A, C, E, G, I, K) embryo at E18.5. Ventral is to the right. Embryos in H and K have also been processed with a bone and cartilage staining procedure. (**L**) A histogram summarizing the observed frequency (X-axis) of the additional phenotypes observed. Note, ‘n=28’ refers to the total number of E17.5 & E18.5 *Prmt5* cKO embryos scored. CTRL embryos did not display any of these phenotypes. Acronyms include hyperpigmentation (HP), syndactyly (SN), kinked tail (KT), no whiskers (NW), and oral blistering (OB). Scale bar = 2mm.

**Supplementary Figure 3:**
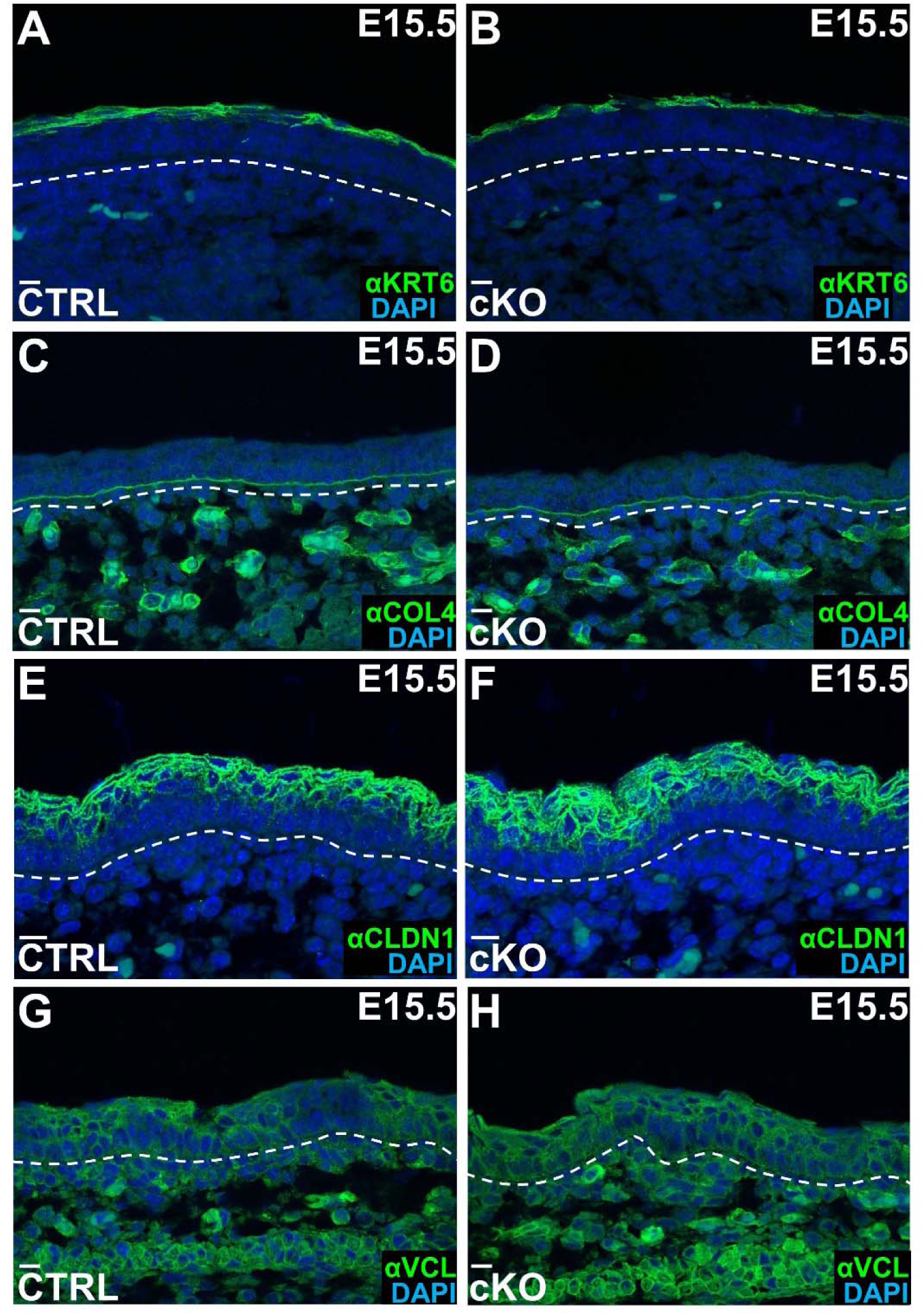
PRMT5-deficient skin retains normal periderm, cell-cell junctions, and basement membrane. (**A-H**) Coronal sections from E15.5 CTRL (A, C, E, G) or *Prmt5* cKO (B, D, F, H) embryonic heads, stained for α-KRT6 (A-B), α-COL4 (C-D), α-CLDN1 (E-F), or α-VCL (G-H) by immunofluorescence. Note, sections shown are the craniofacial epidermis, lateral to the oral cavity, in a coronal plane. White dashed lines demarcate the boundary of the dermis and epidermis. All sectional immunofluorescence images are five Z-stacks compiled with ‘Max Intensity’ in FIJI. Scale bar = 10µm.

**Supplemental Figure 4:**
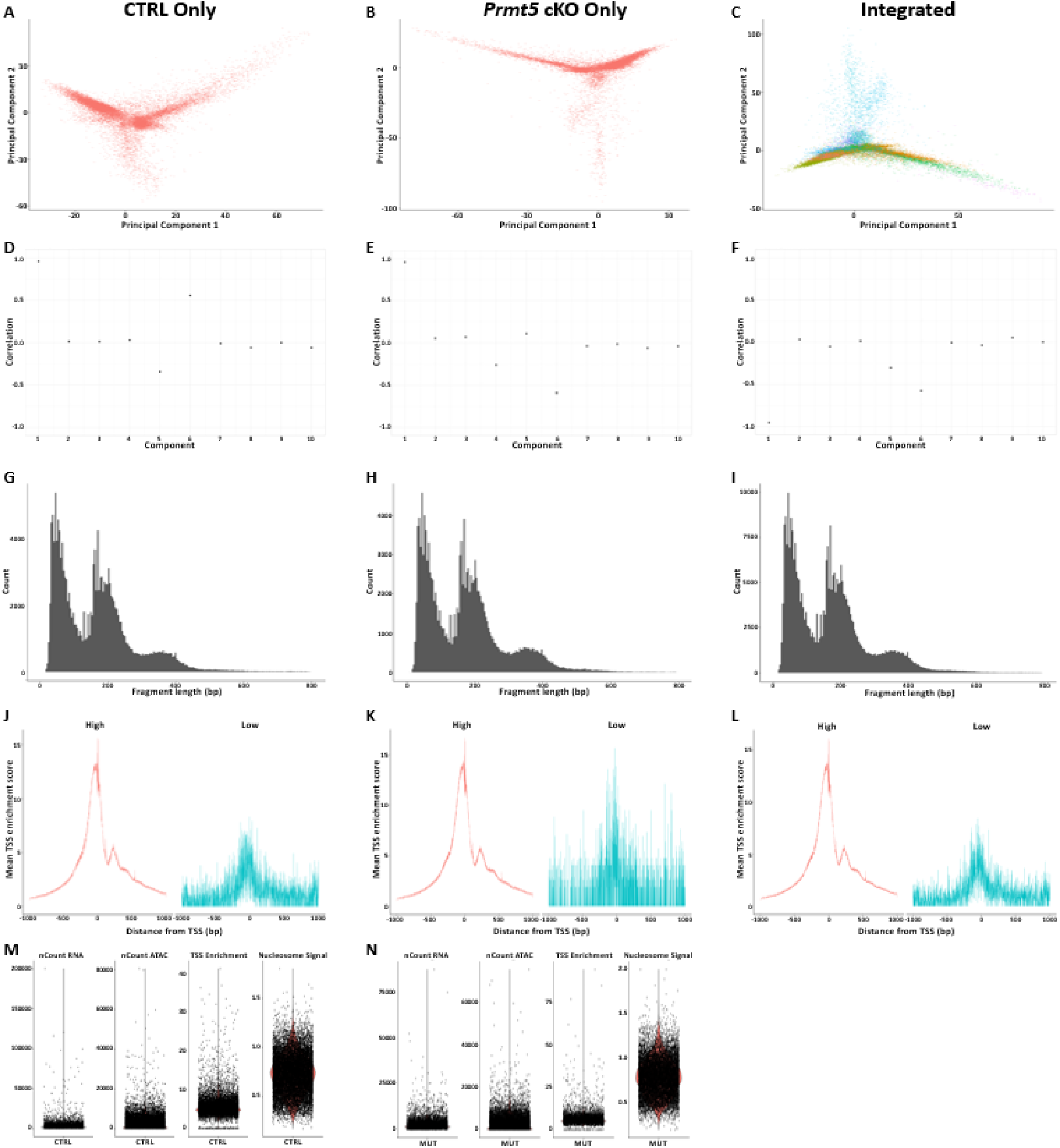
Assessment of quality control metrics confirms the quality of multiome datasets. (**A-C**) Visualization of the first two principal components of CTRL (A), *Prmt5* cKO (B), and integrated (C) datasets using Seurat’s DimPlot function. (**D-F**) Correlation between total ATAC counts and each reduced dimension component of CTRL (D), *Prmt5* cKO (E), and integrated (F) datasets using Signac’s DepthCor function. (**G-I**) Nucleosome signal is visualized using fragment length in CTRL (G), *Prmt5* cKO (H), and integrated (I) datasets using Signac’s Nucleosome Signal function. (**J-L**) Transcription start site enrichment of CTRL (J), *Prmt5* cKO (K), and integrated (L) datasets using Signac’s TSS Enrichment function. (**M-N**) Violin plot of initial nCount RNA, nCount ATAC, TSS Enrichment, and Nucleosome Signal metrics in CTRL (M) and cKO (N) datasets.

**Supplemental Figure 5:**
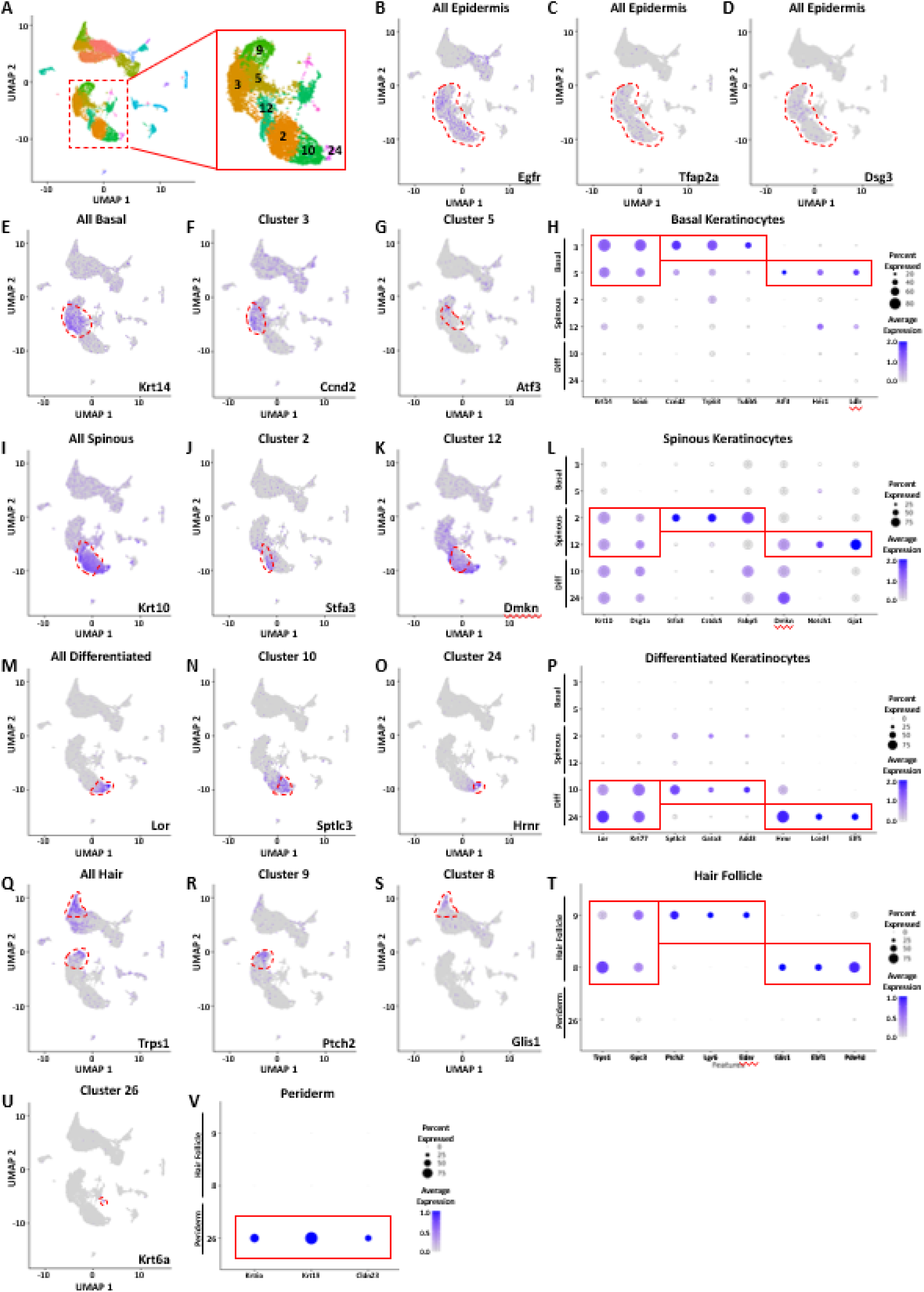
Canonical markers define the keratinocyte, hair follicle, and periderm lineages. (**A**) Labeled UMAP of epidermal derived cells including basal keratinocytes (3 & 5), spinous keratinocytes (2 & 12), differentiated keratinocytes (10 & 24), hair follicle progenitors (9), and periderm cells (26). (**B-D**) Markers of the epidermis enriched in all epidermal clusters. (**E-H**) Markers of basal keratinocytes are shared between Clusters 3 & 5. Whereas markers of earlier differentiation are enriched in Cluster 3 (F, H) and markers of later differentiation are enriched in Cluster 5 (G-H). (**I-L**) Markers of spinous keratinocytes are shared between Clusters 2 & 12. Whereas markers of earlier differentiation are enriched in Cluster 12 (J-L and markers of later differentiation are enriched in Cluster 2 (K-L). (**M-P**) Markers of differentiated keratinocytes are shared between Clusters 10 & 24. Whereas markers of earlier differentiation are enriched in Cluster 10 (N,P) and markers of later differentiation are enriched in Cluster 24 (O-P). (**Q-T**) Markers of the entire hair lineage are enriched in the hair follicle placode (Cluster 9) and the dermal condensate (Cluster 8). Whereas markers of the hair follicle are enriched in Cluster 9 (R,T) and markers of the dermal condensate are enriched in Cluster 8 (S-T). (**U-V**) Markers of the periderm are enriched in Cluster 26.

**Supplemental Figure 6:**
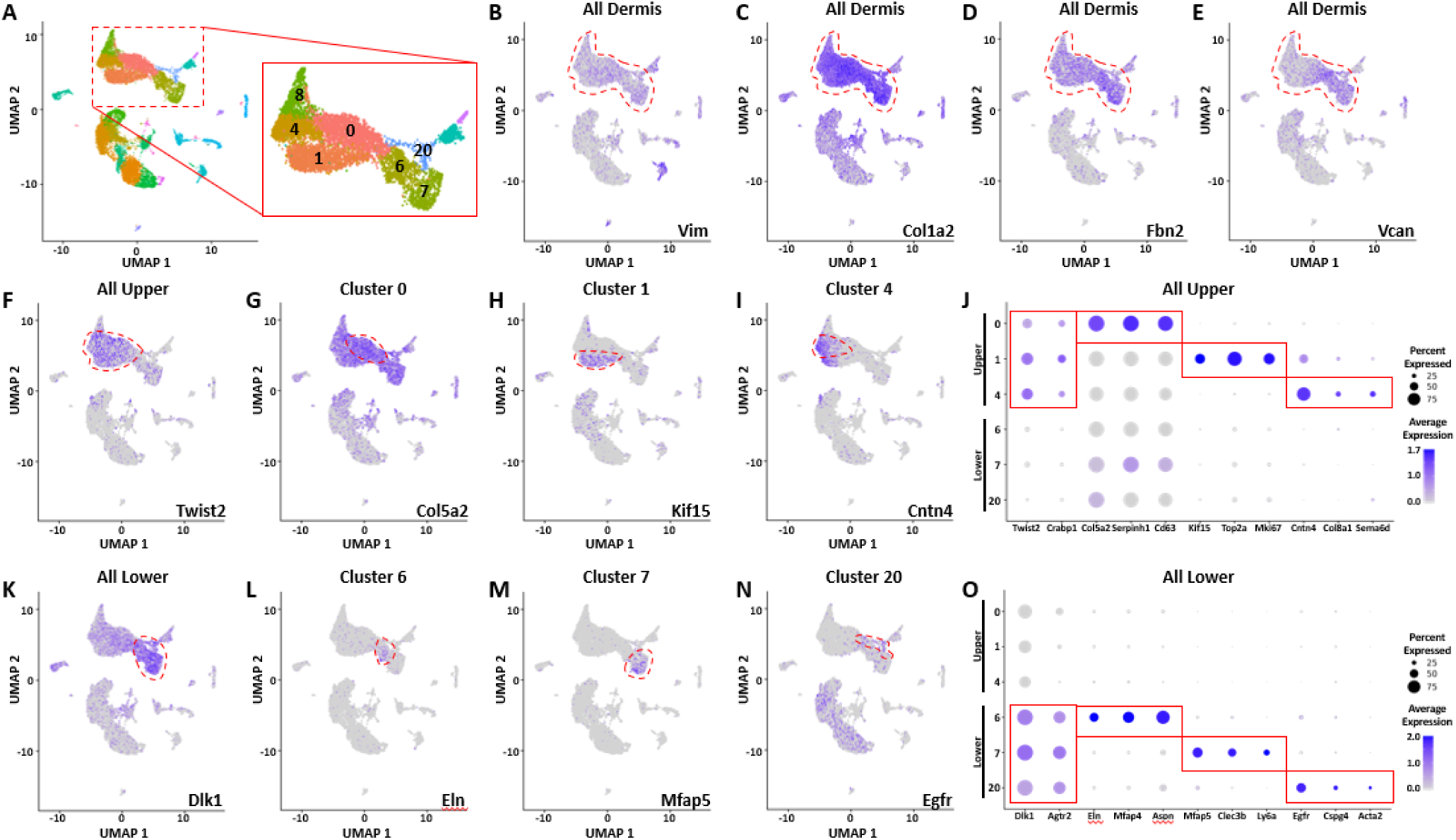
Canonical markers define the upper and lower dermal lineages. (**A**) Labeled UMAP of dermal derived cells including the dermal condensate (8), upper dermis (0, 1, 5) and lower dermis (6, 7, 20). (**B-E**) Markers of the dermis are enriched in all dermal clusters. (**F-J**) Markers of the upper dermis are shared between Clusters 0, 1, and 4. Wherea generic markers of upper dermis are enriched in Cluster 0 (G,J), markers of high levels of proliferation are enriched in Cluster 1 (H, J), and markers of early dermal condensate development are enriched in Cluster 4 (I-J). (**K-O**) Markers of the lower dermis are shared between Clusters 6, 7, and 20. Whereas generic markers of the lower dermis are enriched in Cluster 6 (L,O), markers of fascia fibroblasts are enriched in Cluster 7 (M,O), and markers of mural cells are enriched in Cluster 20 (N-O).

**Supplemental Figure 7:**
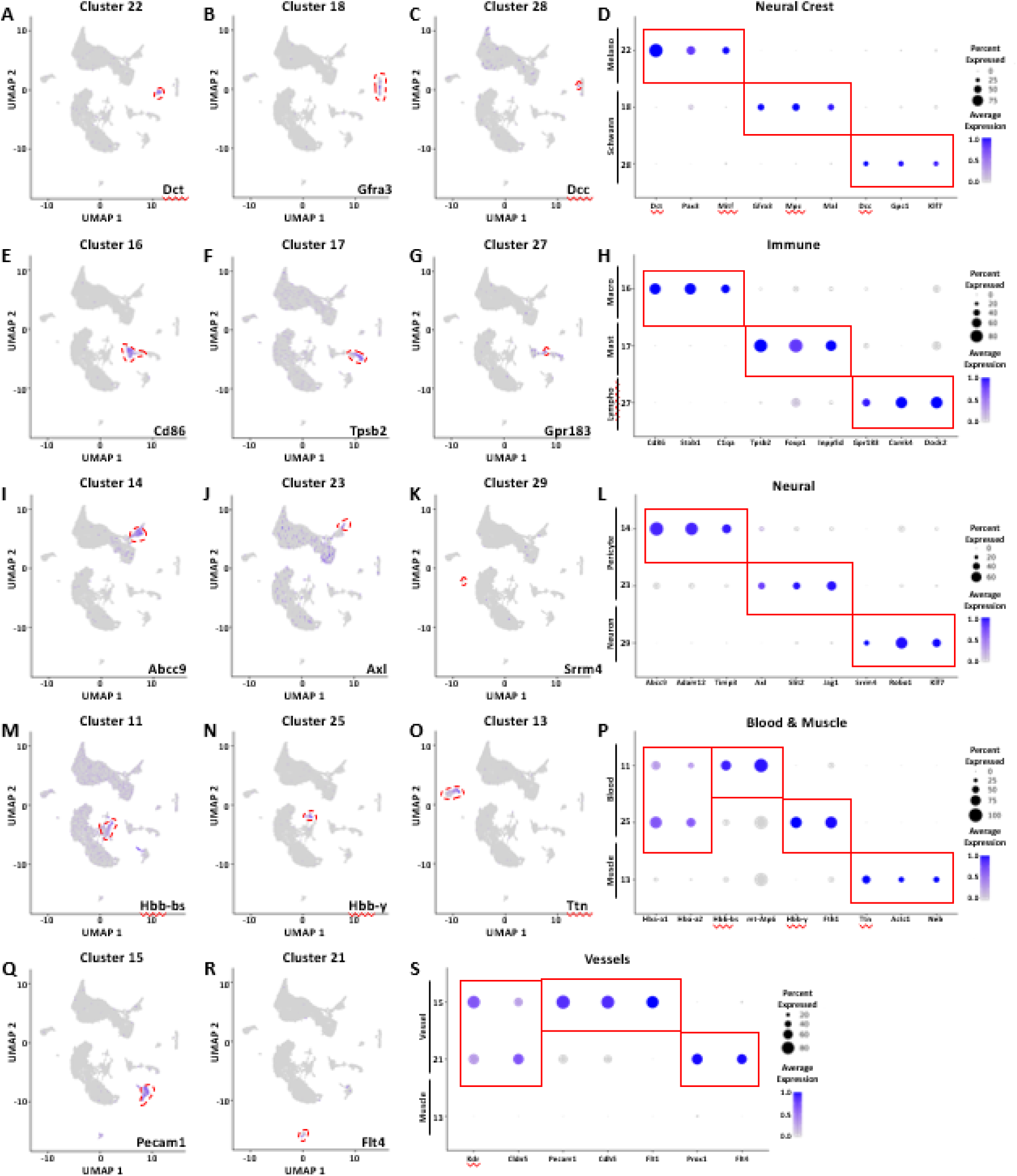
Canonical markers define the neural crest, immune, neural, blood, muscle, and vessel lineages. (**A-D**) Markers of the melanocyte lineage are enriched in Cluster 22 (A,D), markers of the early Schwann cells are enriched in Cluster 18 (B,D), and markers of differentiated Schwann cells are enriched in Cluster 28 (C-D). (**E-H**) Markers of the macrophage lineage are enriched in Cluster 16 (E,H), markers of the mast cell lineage are enriched in Cluster 17 (F,H), and markers of lymphocytes are enriched in Cluster 27 (G-H). (**I-L**) Markers of the early pericyte cells are enriched in Cluster 14 (I,L), markers of differentiated pericyte cells are enriched in Cluster 23 (J,L), and markers of neurons are enriched in Cluster 29 (K-L). (**M-P**) Markers of the blood lineage are shared between Clusters 11 and 25. Whereas markers of mitochondrial identity are enriched in Cluster 11 (M,P) and markers of iron enrichment are enriched in Cluster 25 (N,P). Markers of the muscle lineage are enriched in Cluster 13 (O-P). (**Q-S**) Markers of vessels are shared between Clusters 15 and 21. Whereas markers of vascular endothelial cells are enriched in Cluster 15 (Q,S) and markers of lymphatic endothelial cells are enriched in Cluster 21 (R-S).

**Supplemental Figure 8:**
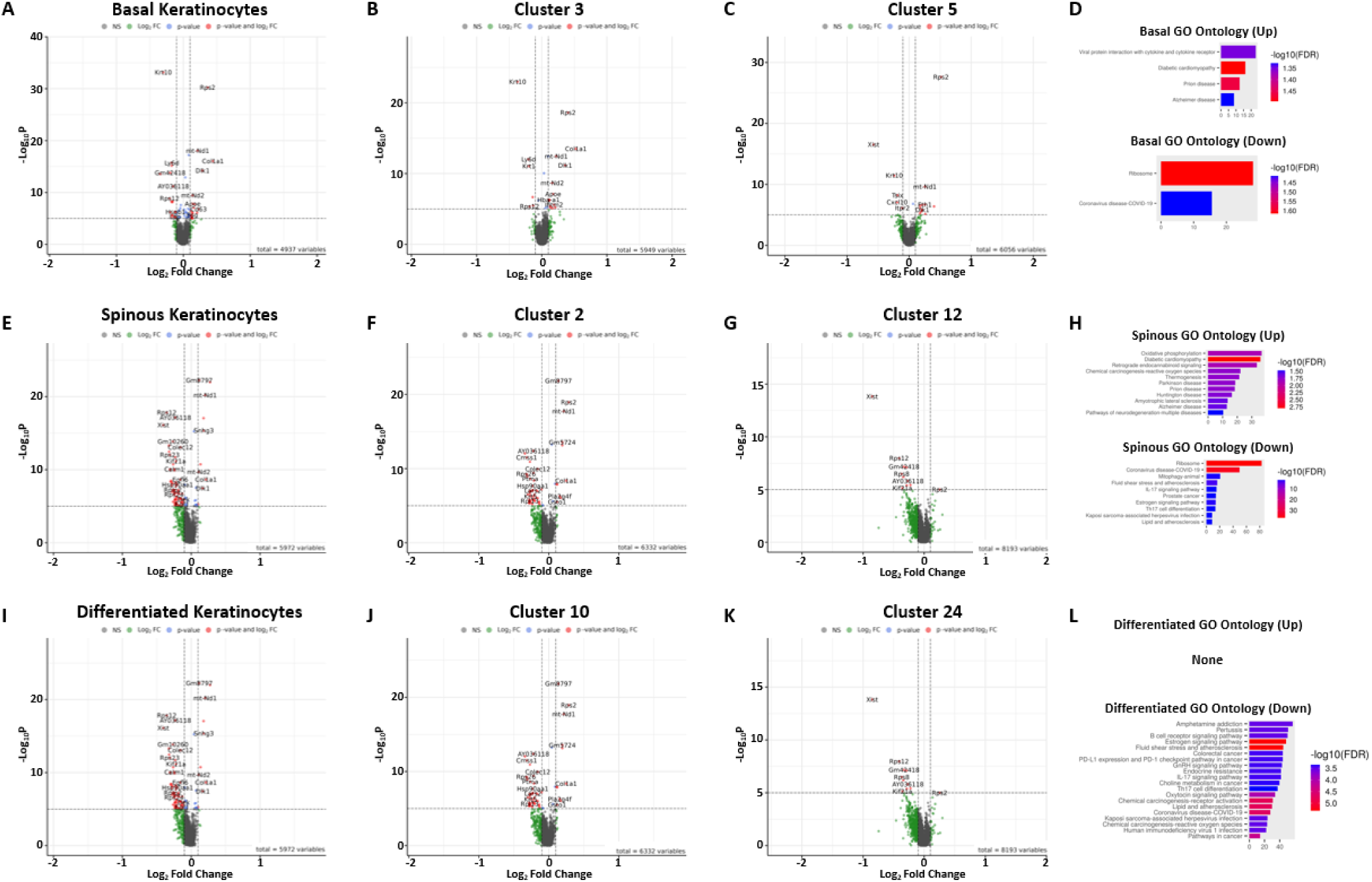
Differential expression between CTRL and *Prmt5* cKO keratinocyte clusters. (**A-D**) Volcano plot (A-C) and GO ontology (D) of differentially expressed genes in all basal clusters (A, D), cluster 3 only (B), and cluster 5 only (C). (**E-H**) Volcano plot (E-G) and GO ontology (H) of differentially expressed genes in all spinous clusters (E, H), cluster 2 only (F), and cluster 12 only (G). (**I-L**) Volcano plot (I-K) and GO ontology (L) of differentially expressed genes in all differentiated clusters (I, L), cluster 10 only (J), and cluster 24 only (K). Differentially expressed genes identified using Seurat v4.3.0 “Find Markers” with a min.pct of 0.1 and logfc.threshold of 0.01.

**Supplemental Figure 9:**
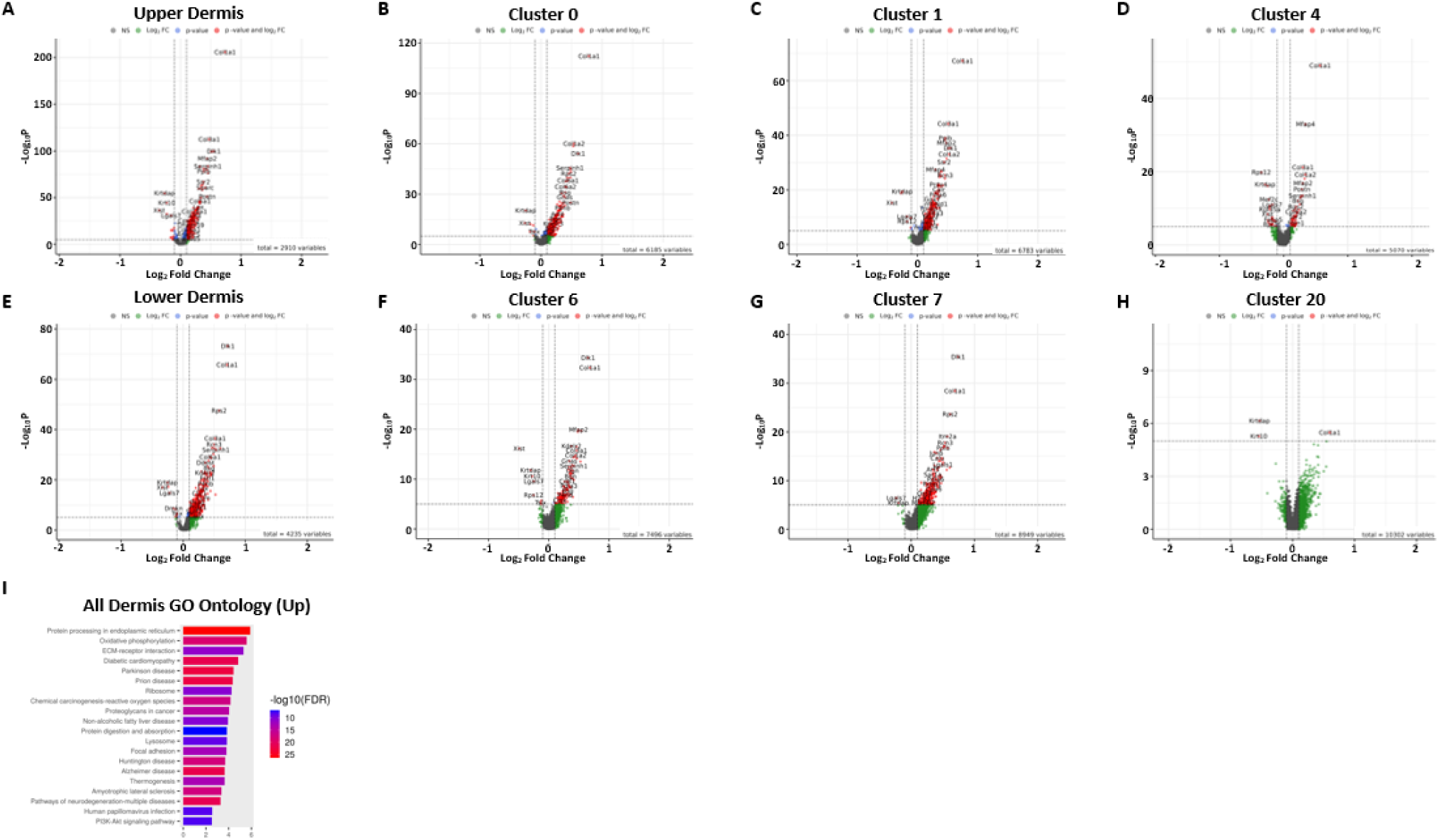
Differential expression between CTRL and *Prmt5* cKO dermal clusters. (**A-D**) Volcano plot of differentially expressed genes in all upper dermis clusters (A), cluster 0 only (B), cluster 1 only (C), and cluster 4 only (D). (**E-F**) Volcano plot of differentially expressed genes in all lower dermis clusters (E), cluster 6 only (F), cluster 7 only (G) and cluster 12 only (H). (**I**) GO Ontology of upregulated genes within all dermal clusters. No GO terms were significantly enriched in the corresponding downregulated genes. Differentially expressed genes identified using Seurat v4.3.0 “Find Markers” with a min.pct of 0.1 and logfc.threshold of 0.01.

**Supplemental Figure 10:**
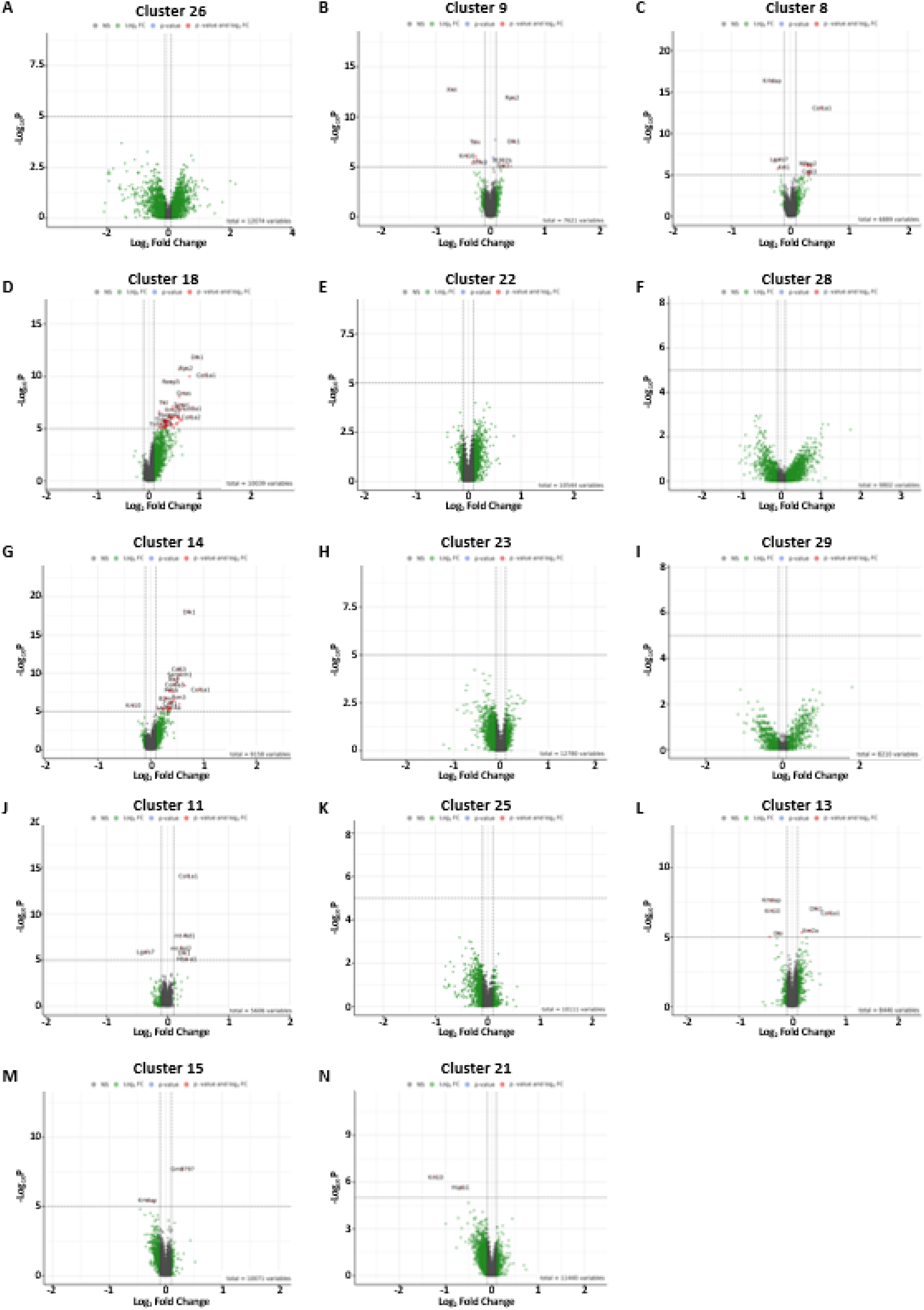
Differential expression between CTRL and *Prmt5* cKO accessory clusters. (**A-N**) Volcano plot of differentially expressed genes in the periderm/cluster 26 (A), hair follicle/cluster 9 (B), dermal condensate/cluster 8 (C), melanocytes/cluster 18 (D), Schwann cells/clusters 22 & 28 (E/F), pericytes/clusters 14 & 23 (G/H), neurons/cluster 29 (I), blood/clusters 11 & 25 (J/K), muscle/cluster 13 (L), and vessels/clusters 15 & 21 (M/N). Differentially expressed genes identified using Seurat v4.3.0 “Find Markers” with a min.pct of 0.1 and logfc.threshold of 0.01.

**Supplemental Figure 11:**
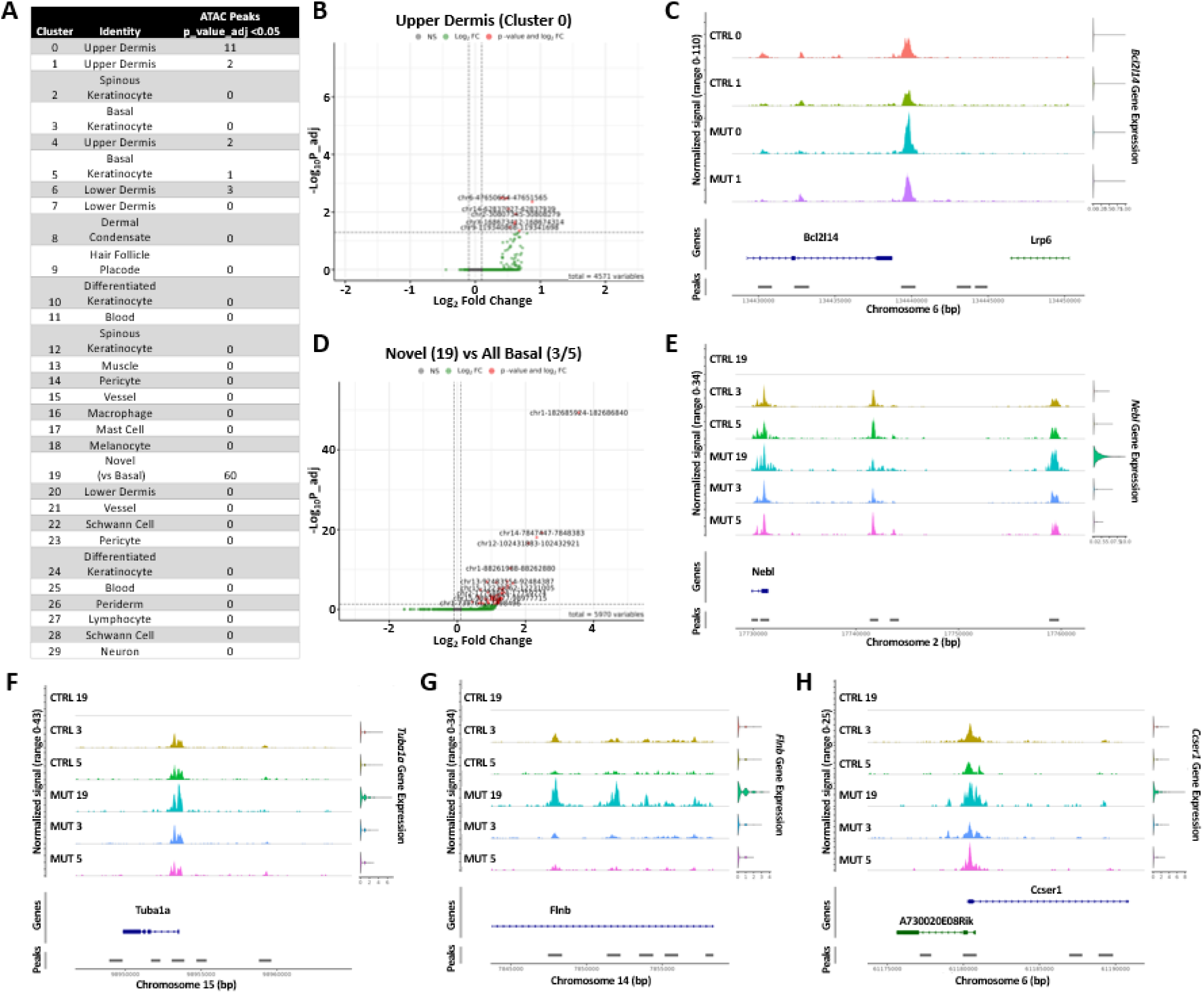
Differential accessibility between CTRL and *Prmt5* cKO clusters. (**A**) Counts of differentially accessible peaks that reach significance (p adj <0.05) in each cluster. (**B**) Volcano plot of differentially accessible peaks in Cluster 0 (Upper Dermis). (**C**) IGV view of the highest fold change peak within Cluster 0 differentially accessible peaks. Violin plot of gene expression is on the right of the IGV view. (**D**) Volcano plot of differentially accessible peaks in Novel (Cluster 19) vs Basal Keratinocytes (Clusters 3 & 5). (**E-H**) IGV views of selected differentially accessible peaks. Violin plots of gene expression are on the right of the IGV view. Differentially accessible peaks identified using Seurat v4.3.0 “Find Markers” with a min.pct of 0.1 and logfc.threshold of 0.01.

**Supplemental Figure 12:**
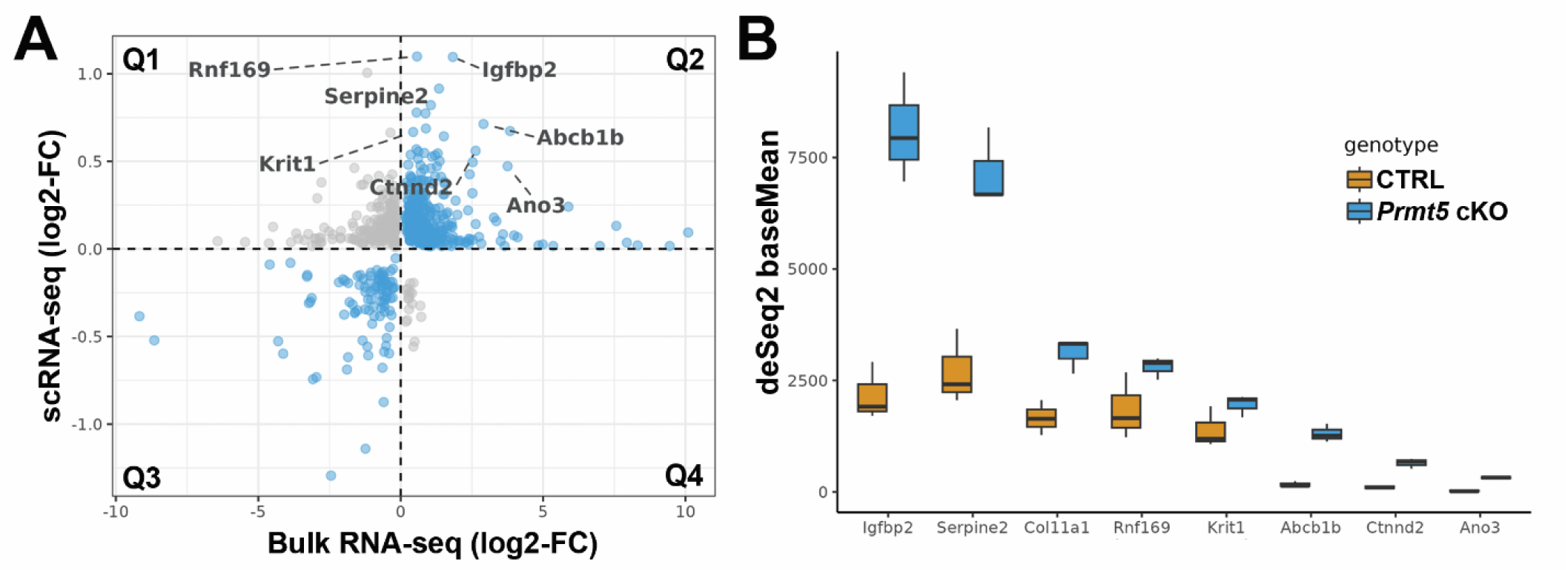
Bulk RNA-sequencing of *Prmt5* cKO and CTRL back skin validates several scRNA-seq ‘novel-cluster’ findings. (**A**) Scatterplot highlighting similar (blue points, quadrant 2 and 3) and divergent (grey points, quadrant 1 and 4) differentially expressed genes identified by bulk RNA-sequencing (X-axis) and scRNA-seq analysis (Y-axis). On the X axis, significant changes in gene expression are plotted as log2 fold change between *Prmt5* cKO and CTRL samples (p_adj < 0.01). On the Y axis, significant changes in gene expression are plotted as log2 fold change between the *Prmt5* cKO-associated novel cluster and the ‘all basal clusters’ (i.e., cluster 3 and 5 from CTRL and *Prmt5* cKO) (p_adj < 0.05). A subset of genes up in both (i.e., Quadrant 2) are labeled and plotted separately in panel B. **(B**) Bar chart highlighting several DEGs significantly upregulated in both bulk RNA-sequencing and scRNA-seq datasets, plotted by gene expression levels (‘baseMean’) in the bulk RNA-sequencing dataset, between genotypes.

**Supplemental Figure 13:**
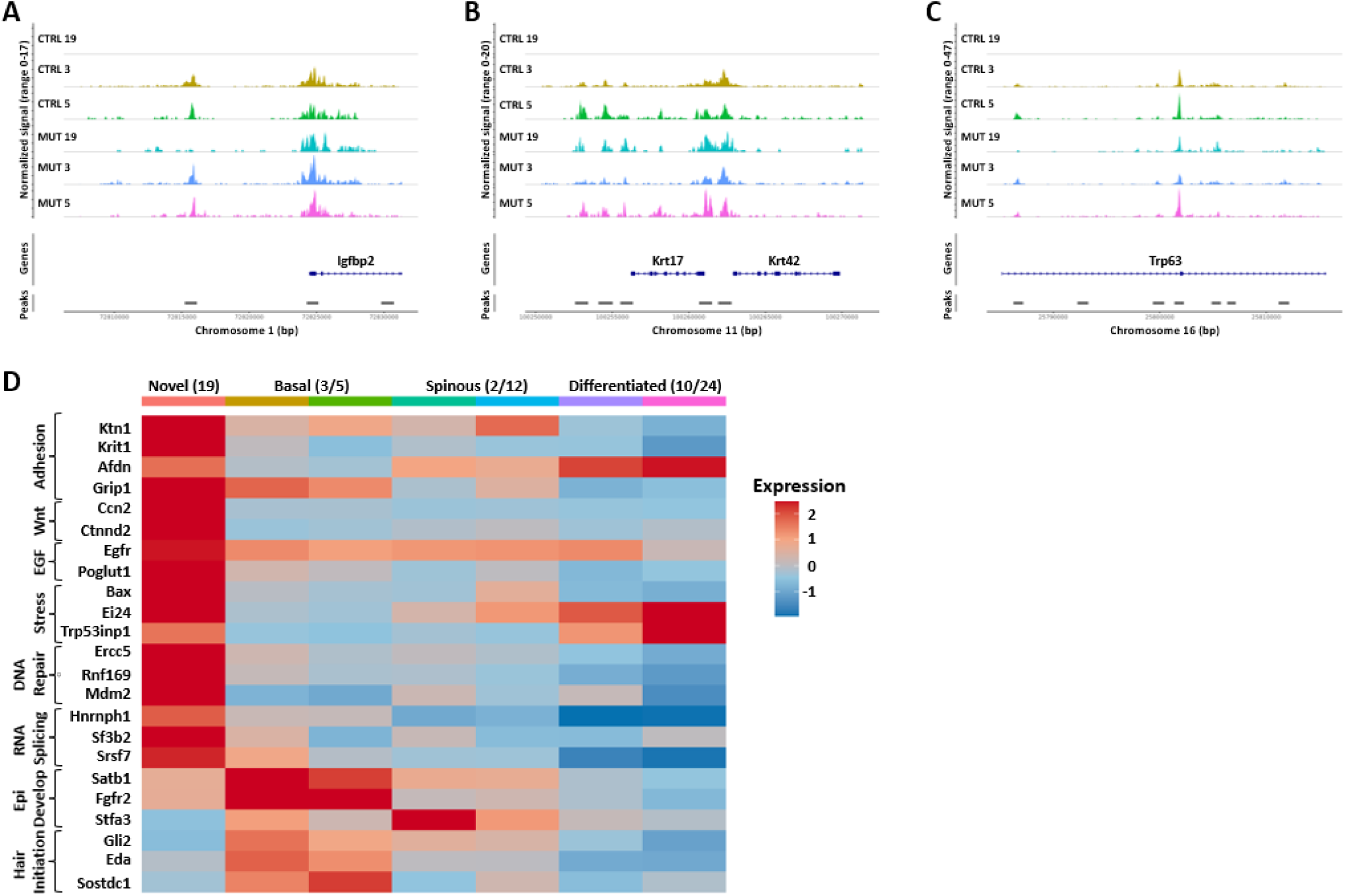
Accessibility analysis of selected genes and additional pathway analyses in the novel cluster. (**A-C**) IGV views of *Igfbp2*, *Krt17*, and *Trp63*, corresponding to their expression changes in Figure 6. (**D**) Heatmap of select genes (rows) and their average expression level (red = high, blue = low) in distinct epidermal clusters (rows). Each cluster of genes is sectioned into their corresponding broad category.

**Supplemental Table 1: Cluster RNA Markers**

**Supplemental Table 2: Differential Expression**

**Supplemental Table 3: Differential Accessibility**

